# A ganglioside-based senescence-associated immune checkpoint

**DOI:** 10.1101/2021.04.23.440408

**Authors:** Charlène Iltis, Laetitia Seguin, Ludovic Cervera, Lou Duret, Tynhinane Hamidouche, Sarah Kunz, Olivier Croce, Clément Delannoy, Yann Gueŕardel, Fabrice Allain, Lyvia Moudombi, Paul Hofman, Estelle Cosson, Julien Guglielmi, Thierry Pourcher, Véronique M. Braud, Marina Shkreli, Marie-Cécile Michallet, Chloé C. Feral, Eric Gilson, Julien Cherfils-Vicini

## Abstract

Senescent cells accumulate in aging tissues, and their elimination can favor healthy aging^1-4^. Therefore, therapeutic interventions targeting cellular senescence may be promising strategies for delaying or reversing a vast range of age-related diseases^5^. As cells of the immune system are responsible for senescent cell elimination^6-11^, a possible anti-aging and pro-healthspan treatment is the specific activation of the immune system to induce senescent cell clearance. However, whether this elimination is limited by an immune checkpoint leading to tolerance of senescence cells is currently unknown. Here, we show that cellular senescence, elicited by various stressors other than oncogenic activation, triggers immune escape toward natural killer (NK) cells, which may thus limit the use of anti-senescence immunotherapies. Moreover, using mass spectrometry, we reveal that senescent cells reshuffle their glycosphingosine composition, toward a marked increase in the ganglioside content, including the appearance of disialylated ganglioside GD3. This senescence associated GD3 overexpression results from transcriptional upregulation of the gene encoding the enzyme ST8SIA1, which is responsible for GD3 synthesis. The high level of GD3 leads to a strong immunosuppressive signal affecting NK cell-mediated immunosurveillance. In a mouse model of lung fibrosis, senescent cell-dependent NK cell immunosuppression is blunted by *in vivo* administration of anti-GD3 monoclonal antibodies leading to a clear anti-fibrotic effect. These results demonstrate that GD3 upregulation in senescent cells drives a switch from immune clearance toward immune tolerance of senescent cells. Therefore, we propose that GD3 level acts as a senescence-associated immune checkpoint (SIC) that regulates NK cell functions toward senescent cells. Thus, targeting GD3 with specific antibodies may be a promising strategy for the development of effective anti-senescence immunotherapies.

---

Aging is a process involving loss of cell and tissue functionality that is mediated in part by the accumulation of senescent cells, but the factors regulating this accumulation remain largely unknown. The immune system plays a major role in the senescent cell elimination, essentially through Natural Killer (NK) cells, macrophages and CD4+ T cells^6,7,9,12,13^ acquiring senescence-associated secretory phenotype (SASP). Besides, senescent cells can also be immunosuppressive^8,11^. Therefore, a better understanding of senescence and immune system interplay may lead to new therapeutical options for age-related disease. To determine the features of senescence immunosurveillance, we performed an *in vivo* Matrigel plug assay, measuring the immune recruitment induced by human replicative senescent cells (Fig.1a). We cultured MRC-5 human lung primary fibroblasts under 5% O^2^ conditions until replicative senescence, defined as at least 90% double Senescence-Associated beta-galactosidase-positive (SA-β-Gal+), 5-ethynyl-2′-deoxyuridine-negative (EdU-) cells (Extended Data Fig. 1a-b) as well as increased DNA damage at telomeres (Extended Data Fig. 1c-e). Consistent with previous reports^8,9,11,13^, the immune recruitment induced by human replicative senescent cells is extensive, with increased infiltration of NK cells (CD3-NKp46+) and neutrophiles (CD11b+ GR1 hi) (Fig. 1b, Extended Data 2a-b). Surprisingly, the degranulation of NK cells recruited by senescent cells is reduced three-fold compared to NK cells recruited by young cells (Fig.1c). In an *in vitro* co-culture experiment with mouse NK cells (Fig. 1d), we observed that senescent cells inhibit more effectively than their young counterparts NK cell degranulation, but not interferon gamma (IFN-γ) production (Fig. 1e; Extended Data 2f-g). Beyond degranulation inhibition, replicative senescent MRC5 inhibit also human NK cell specific killing (Fig.1f). The SASP of the senescent cells did not alter immune infiltration and had no effects on NK cell recruitment or function (Extended Data 2c-e).

**Figure 1:**
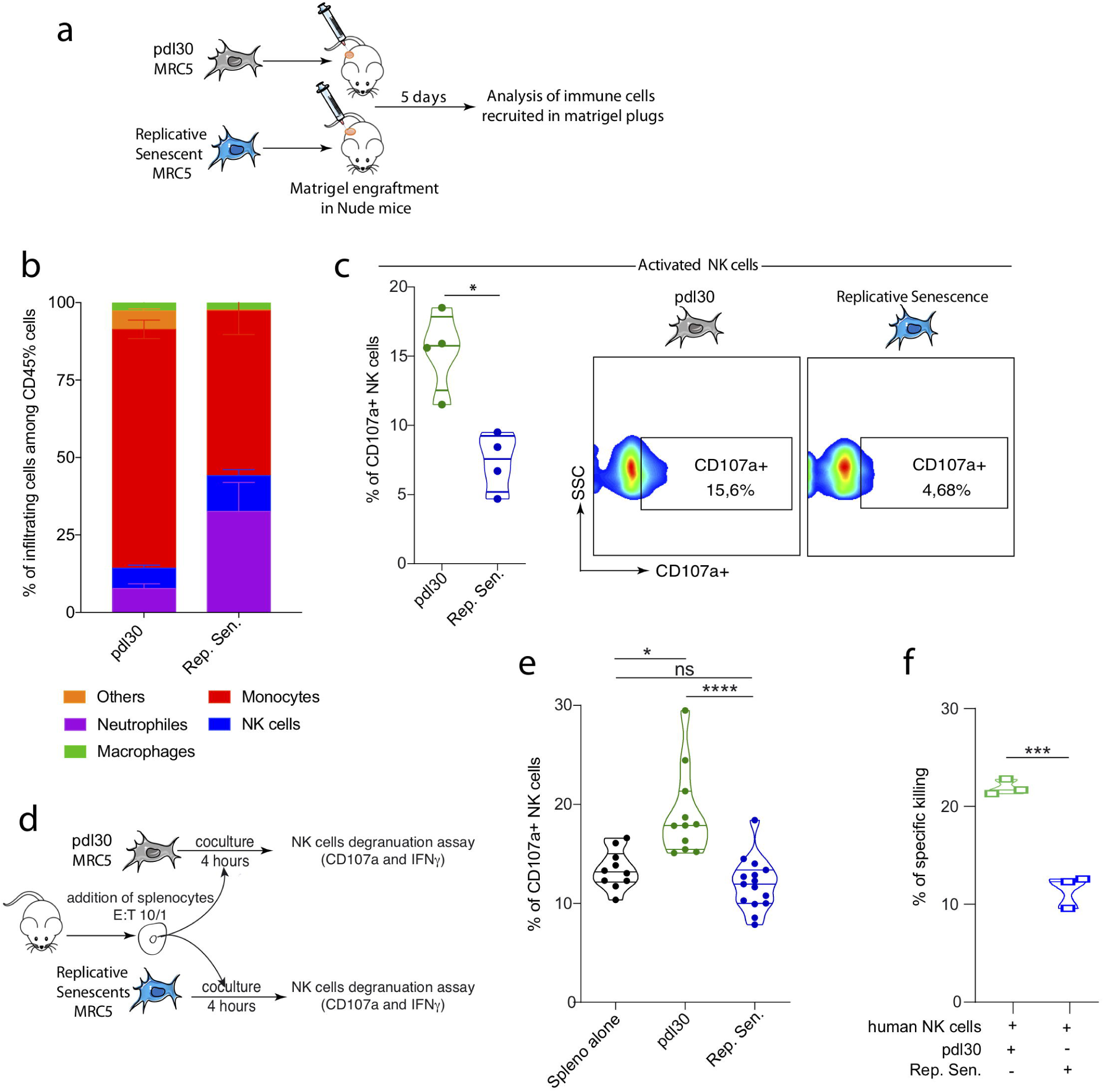
Human replicative senescent cells recruit NK cells *in vivo* but inhibit their degranulation in a SASP independent mechanism. **a**, scheme of the Matrigel plug assay. **b**, quantification of the immune cell infiltration induced by human replicative senescent MRC5 cells or proliferative MRC5 cells after 30 population doubling level (pdl30). **c**, Quantification of the NK cell degranulation within the Matrigel plug assay. **d**, representative scheme of the *in vitro* coculture assay of human replicative MRC5 senescent cells with mouse splenocytes. **e**, quantification of NK cell degranulation during *in vitro* coculture experiments. **f**, *in vitro* killing assay of pdl30 MRC5 or replicative senescent MRC5 by primary human purified NK cells. All experiments are performed with n = 4 mice per group; *p < 0.05, **p < 0.01, and ***p < 0.001; Mann–Whitney *U* test (1a-c) or represent the mean of n = 3 independent experiments; *p < 0.05, **p < 0.01, and ***p < 0.001; Student’s t-test (**1d-f**).

Because SASP appeared insufficient to mimic the effects of senescent cells on immune infiltration and NK cell inhibition (Extended Data 2c-e), we hypothesized that cell surface molecules may be involved. We analyzed the glycocalyx composition of proliferative and senescent MRC5 cells using mass spectrometry, focusing on glycosphingolipids (Fig. 2a), O-glycans (Extended Data 3a-c) and N-glycans (Extended Data 4a-b). Mass spectrometry analysis showed that MRC5 *N*-glycome was made of oligomannosylated (Man_5_GlcNAc_2_ to Man_9_GlcNAc_2_), neutral and complex sialylated LacNAc containing *N*-glycans, whereas *O*-glycome was exclusively constituted of mono and disialylated short *O*-glycans (Extended Data 3b and 4a). *N*-glycome and *O*-glycomes were not significantly modified in senescent MRC5 compared to control MRC5 cells (Extended Data 3c and 4b). In contrast, the glycosphingolipids pattern of senescent MRC5 cells was dramatically modified compared to MRC5 control cell. While young cells contained both neutral globosides (Gb3 and Gb4/GA1) and gangliosides (GM3, GM2 and GM1 sialylated), replicative senescent cells contained primarily gangliosides with a marked upregulation of disialylated gangliosides (GD3) (Fig. 2a). Flow cytometry and immunofluorescence analysis (Fig. 2b-c; Extended data 5a) confirmed the higher level of GD3 in replicative senescent cells. To decipher the mechanism, we measured the transcription of enzymes involved in ganglioside biosynthesis using quantitative polymerase chain reaction (RTqPCR) (Extended Data 5b). ST8SIA1, the enzyme that generates GD3 by adding sialic acids, increases sharply in replicative senescent cells (Extended Data 5c). ST8SIA1 expression does not increase progressively during the cell proliferative history, but, instead, is expressed massively at the onset of senescence (Extended Data 5d). Moreover, GD3 is not cleaved or secreted, as we were unable to detect it in the supernatant of proliferative MRC5 or replicative senescent MRC5 cells (Extended Data 5e). Such upregulation of *ST8SIA1* gene expression together with the elevated level of GD3 were observed in other contexts associated with premature induction of senescence in response to a wide range of stressors (Extended Data 5f-h). In contrast, *ST8SIA1* and GD3 expression is downregulated in senescent cells induced by an oncogenic stress (Extended Data 6a-d). Accordingly, in the lungs of 2-month-old KRasG12D-expressing mice, oncogene-induced senescence (OIS) cells, defined as SA-β-Gal+ cells, do not express GD3 (Extended Data 6e-f).

**Figure 2:**
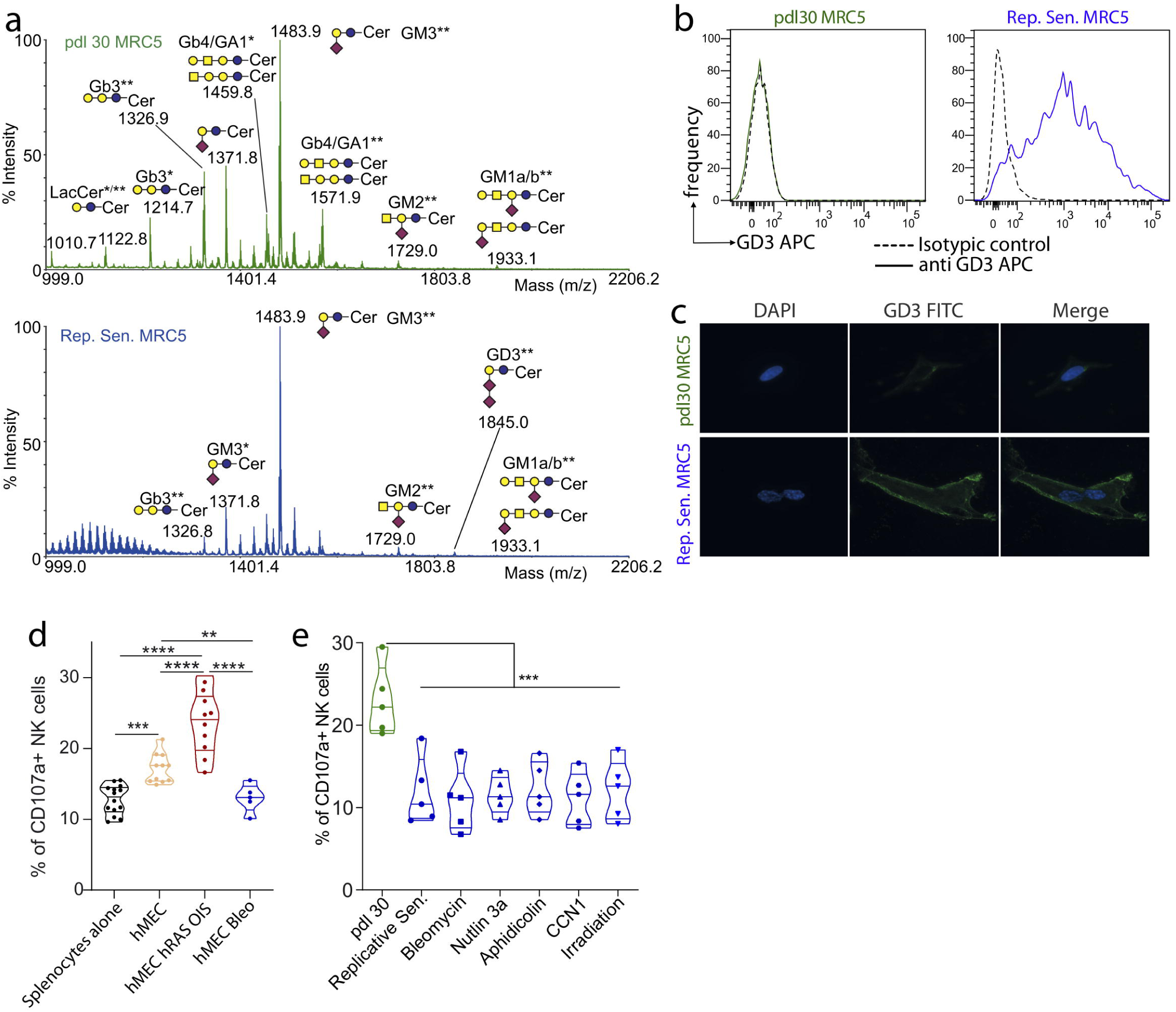
The ganglioside GD3 is strongly expressed by Senescent cells and inversely correlates with their immunogenic properties towards NK cells. **a**, mass spectrometry analysis of permethylated glycosphingolipds from human young or replicative senescent MRC5 cells; GSLs are present as d18:1/C16:0 (*) and d18:1/C24:0 (**) isomers. **b-c**, analysis of GD3 expression in human young or replicative senescent MRC5 cells, either by FACS (**b**) or Immunofluorescence (**c). d-e**, *in vitro* co-culture experiment with Oncogene or bleomycin induced senescence in hMEC cells (**d**) or young, replicative or stress induced senescent MRC5 cells (**e**) with mouse splenocytes. Data represent the mean of n = 3 independent experiments, *p < 0.05, **p < 0.01, and ***p < 0.001; Mann–Whitney *U* test.

As GD3 triggers the inhibitory immunoreceptor tyrosine-based inhibitory motif (ITIM) receptor Siglec-7 (or Siglec-E/H in mice)^14,15^, the interaction GD3 (senescent cells) / Siglec-7 (NK cells) could mediate the observed NK cell inhibition caused by senescent cells. Therefore, we analyzed the impact of senescent cells that either express or repress GD3 on NK cell functions *in vitro*. While OIS cells that do not express GD3 can increase NK cell degranulation (Fig. 2d), all the other types of senescent cells expressing GD3 abolished NK cell degranulation (Fig. 2e). To directly assess whether GD3 expression by senescent cells regulates their immunogenic potential toward NK cells, we modulated the GD3 level in senescent cells through enzymatic treatment (neuraminidase which hydrolyzes terminal N- or O-acyl neuraminic acids; Extended Data 7a), *ST8SIA1* knock-down (KD; Extended Data 7b-d) or ST8SIA1 overexpression in young cells (Extended Data 7e-g) before exposing them to murine NK cells in a co-culture assay (Fig. 3a). GD3 inhibition using neuraminidase or via KD of *ST8SIA1* totally rescued the capacity of senescent cells to activate NK cell degranulation (Fig. 3b-c) but did not increase IFN-γ production (Extended Data 7h). Conversely, overexpression of *ST8SIA1* by young dividing cells (Extended data 7e-g) inhibited mouse NK cell degranulation which can be inhibited by neuraminidase treatment (Fig. 3d). As GD3 is a cell surface molecule that can be targeted with antibodies, we explored whether NK cell degranulation could be restored using an anti-GD3 monoclonal antibody (Fig. 3e; Extended Data 8). GD3 targeting by the monoclonal antibody was sufficient to restore NK cell degranulation and killing. This NK cell functionality rescue by anti-GD3 mAb is achieved in a dose-dependent manner, achieving full rescue with 2 µg of antibody (Fig. 3e), probably by inhibiting the interaction between GD3 and the inhibitory receptor Siglec-E/H as well as by enhancing antibody-dependent cellular cytotoxicity (ADCC)^16^. Importantly, GD3 expression is also sufficient to inhibit killing of senescent cells by human primary NK cells (Extended Data 8e). The inhibition of human senescent cell killing by NK cells is GD3-dependent, as revealed by rescue of the killing phenotype after anti-GD3 monoclonal antibody (mAb) treatment (Extended Data 8e). Together, these results demonstrate that GD3 expression by senescent cells determines their capacity to inhibit NK cells and thus escape elimination by NK cells.

**Figure 3:**
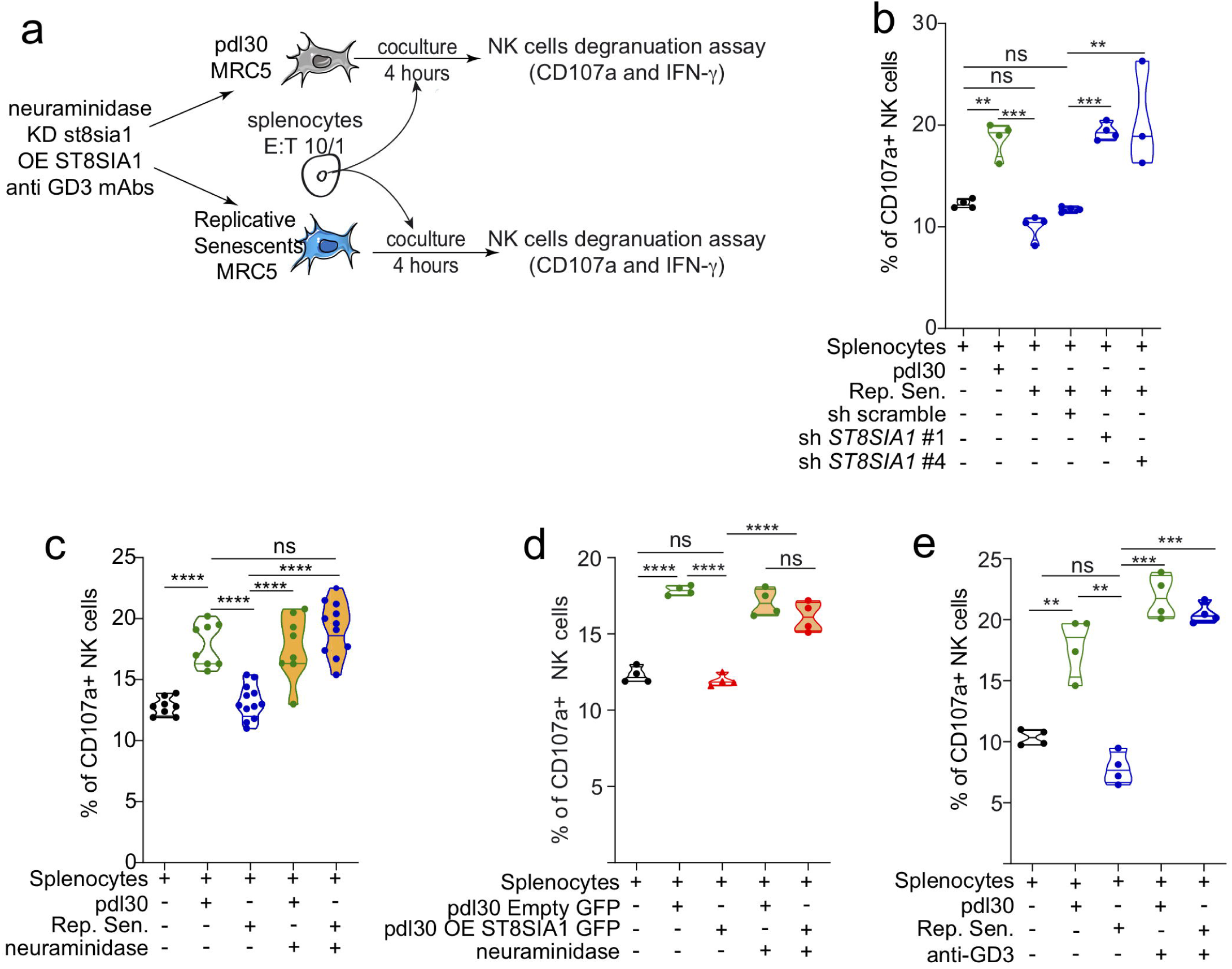
The natural GD3 expression by senescent cells directly determine their NK cell mediated immune surveillance. **a**, Representative scheme of the *in vitro* co-culture experiment. **b-e**, *in vitro* co-culture experiment using either replicative senescent cells knock-down for *ST8SIA1 (***b)** or treated with neuraminidase (**c)**, young cells overexpressing ST8SIA1 and treated or not with neuraminidase (**d**) or replicative senescent cells in presence of anti-GD3 monoclonal antibody or isotype control(**e**). Data represent the mean of n = 4 independent experiments, *p < 0.05, **p < 0.01, and ***p < 0.001; Mann–Whitney *U* test.

To assess the physiological impacts of NK cell surveillance escape by GD3-expressing senescent cells, we analyzed the expression of GD3 by senescent cells in lung fibrosis. Indeed, recent studies have established a clear link between senescent cell accumulation and lung fibrosis^9,17,18^ which can be mimicked in mice through intratracheal instillation of bleomycin^19^ (Fig. 4a). As expected, bleomycin treatment increased collagen deposition (Extended data 9a-b). The fibrotic areas contain SA-β-Gal+ cells expressing GD3 (Fig. 4b-c and Extended Data 9c). Using imaging flow cytometry, we observed that both immune cells (CD45+ cells) end epithelial cells (EpCAM+ CD45-cells) are SA-β-Gal+ (Fig. 4b), with a 4.7-fold-increase in SA-β-Gal+ cell frequency among EpCAM positive cells (Fig. 4c). In contrast to human lung fibrosis, bleomycin-induced fibrosis was shown to partially regress spontaneously^20^. Accordingly, we observed that the fibrotic lesion partially regressed at day 120 (Extended Data 9d), with a significant decrease in collagen deposition (Extended Data 9e, left panel). However, GD3-positive cells remained present 120 days after injury in the remaining fibrotic lesions, and the intensity of GD3 expression was not significantly reduced compared to that on day 21 (Extended Data 9e, right panel). Thus, senescent cells were not eliminated and persisted in the tissues within fibrotic sequelae. As GD3+ senescent cells are found in fibrotic lungs, we tested whether immune infiltration was affected. Through flow cytometry analyses of fibrotic lungs, we revealed that global immune infiltration is slightly altered with an increased frequency of NK cells (Extended Data 9f). To assesses their function, lung infiltrating cells, containing primary NK cells, were incubated *ex vivo* with the NK cell target YAC-1 (Fig. 4d). By flow cytometry, we observed that NK cells from fibrotic lungs degranulated less against YAC-1 cells than their counterparts treated with phosphate-buffered saline (PBS). Moreover, the function of splenic NK cells from the same mice was assessed simultaneously but did not show different activities (Extended Data 9g). Away from the lungs, NK cell function is similar between the control and treatment groups, suggesting that the presence of GD3-positive senescent cells within lung is sufficient to locally inhibit NK cell degranulation *ex vivo*, but not to trigger systemic NK cell inhibition.

**Figure 4:**
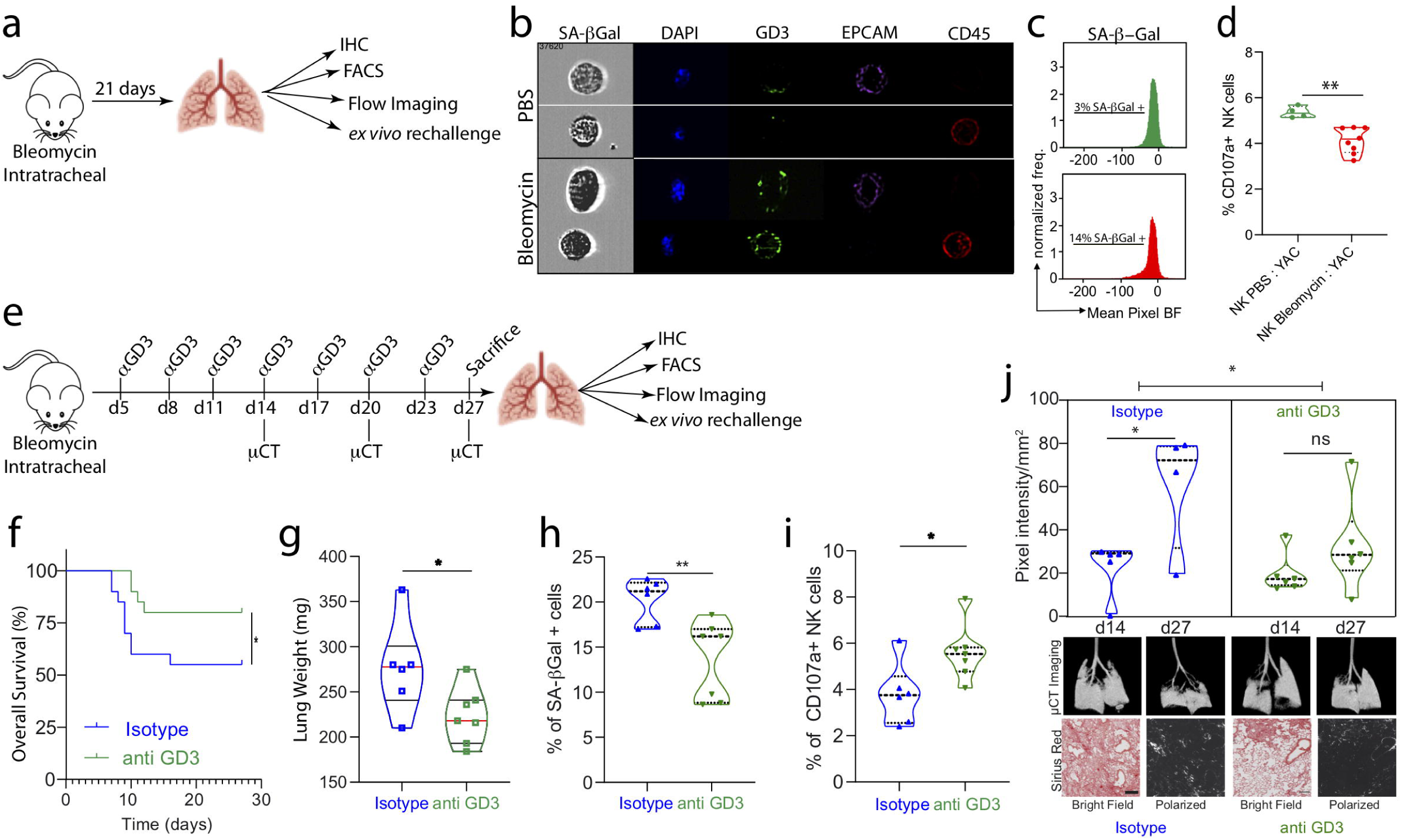
GD3 targeting *in vivo* increases overall mouse survival and reduces lung fibrosis lesions by restoring locally NK cell mediated immune surveillance. **a**, representative scheme of the *in vivo* lung fibrosis experiment. **b-c**, characterization by ImagestreamX of senescent cells using SA-β−Galactosidase assay, GD3, EPCAM and CD45 staining. **d**, Determination of the intrapulmonary NK cell functionality *ex vivo* against YAC-1 cells after 4 hours of rechallenge. **e**, Representative scheme of the *in vivo* anti GD3 mAb treatment in lung fibrosis model. **f**, overall survival analysis of fibrotic mice depending on mAb treatment. **g**, lungs weights after treatment at d27. **h**, quantification of senescent cells infiltrating lungs at d27. **i**, evaluation by flow cytometry of intrapulmonary NK cells functionality at d27. **j**, evaluation of lung fibrosis by µCT imaging at d14 and d27 depending on the treatment. µCT imaging at d14 and d27 and Sirius red staining at d27 for the same mice are presented. All experiments are performed with n = 8 mice per group; *p < 0.05, **p < 0.01, and ***p < 0.001; Mann–Whitney *U* test.

Next, we investigated whether *in vivo* GD3 targeting using a blocking mAb can inhibit the tolerogenic effects of senescent cells locally. Thus, we analyzed the impact of *in vivo* GD3 targeting by monitoring the overall survival and lung fibrotic lesion formation with microcomputed tomography (μCT) imaging, the quantity of senescent cells with imaging flow cytometry, and the immune capacity by flow cytometry (Fig. 4e). Treatment with anti-GD3 significantly improved overall survival (Fig. 4f) and reduced lung fibrosis, as determined by lung weight (Fig. 4g) and lung density measurement via µCT (Fig. 4j). Follow-up analyses of lung fibrosis over time in the same mice revealed that anti-GD3 treatment was sufficient to block disease progression, and for some mice, to reverse it (Fig. 4j upper panel). This result was confirmed through histological analyses of lung fibrosis at the end of treatment and comparison with µCT imaging (Fig. 4j, lower panel). The anti-fibrotic effects of GD3 mAb are associated with a 25% decrease in senescent cell infiltration (Fig. 4h), despite similar GD3 expression in the remaining senescent cells (Ext. Data 10a-b). Although anti-GD3 treatment did not affect the overall proportion of immune cells (Extended Data 10c), it was sufficient to restore NK cell degranulation (Fig. 4i) and activation (Extended Data 10d). In addition, we assessed the functional capacities of lung and spleen NK cells *ex vivo* through a YAC-1 re-challenge experiment (Extended Data 10e-h) where lung cells or splenocytes were co-cultivated with YAC-1 cells. While the NK cell proportion in the lung (Extended Data 10e) or spleen (Extended Data 10g) was not affected by the treatment, lung NK cell function (degranulation and IFN-γ production) against YAC-1 was strongly enhanced *ex vivo* (Extended Data 10f). In contrast, anti-GD3 mAb treatment did not alter spleen NK cell function (Extended Data 10h), showing that NK cell inhibition is local and dependent on the presence of GD3. Altogether, these results showed that the pro-fibrotic effects of senescent cells in the lung are GD3-dependent and favor the inhibition of NK cell-mediated immunosurveillance, which can be restored through anti-GD3 mAb.

Such an association of GD3 expression to fibrosis-associated senescent cells is not restricted to the lung since we observed that senescent cells, defined by SA-β-Galactosidase assay in experimental mouse kidney fibrosis induced by Adriamycin treatment, are GD3-positive (Extended Data 11a). GD3 is strongly expressed in kidney glomeruli after the kidney injury (Extended Data 11b) and the percentage of kidney area (Extended Data 11c, left panel) or glomeruli area (Extended Data 11c, right panel) is strongly increased, revealing that GD3 positive senescent cells following injury is not restricted to damaged lungs. Upon natural aging, we observed that GD3 is not found in a secreted or cleaved form in the serum of old mice (Extended Data 12a) but significantly expressed by senescent cells that naturally appears in old kidney (Extended Data 12b) and that GD3 senescent cells occurrence increases strongly with aging (Extended Data 12c). To analyze the expression of *ST8SIA1* upon human aging, we took advantage of the RNAseq public data set of human samples available through the GTEX consortium. We downloaded data from GTEX and generated a script on order to classify gene expression (in Transcript Per Million) as a function of the age of healthy donors (Extended Data 13). We observed a significant increase of *ST8SIA1* expression in the human lungs with age. Concomitantly, the expression of marker genes is indicative of an increased senescence rate : *p16, p21 or SERPIN E1* gene expression significantly increased while the one of *LMNB1* decreased in the lungs from same donors (Extended Data 13a-b). Importantly, the expressions of *p16* or *p21* are significantly correlated to the expression of *ST8SIA1* (Pearson correlation, p<0,0001; Extended Data 13c-d). These analyses strongly suggest that upon human aging, the senescent cells that accumulate within human lungs are *ST8SIA1* positives.

In summary, here, we unveiled that human senescent cells exhibit strong immunosuppressive functions *in vivo*, with the ability to recruit NK cells and fine-tune their inactivation through a GD3-dependent pathway. This result may explain why some senescent cells are not eliminated from tissues, and thus can accumulate with age. Consequently, our results show that senescent cells alter their cell surface glycolipid repertoires, which confers tolerogenic ability, and thereby modify their interactions with immune cells. Of note, OIS cells behave in a seemingly opposite way, downregulating ST8SIA1/GD3 expression, which is consistent with their capacity to be eliminated by NK cells. Moreover, we show in a model of severe lung fibrosis that GD3 targeting is sufficient to strongly reduce lung fibrosis and improve overall survival by blunting senescent cell-dependent NK cell inhibition (Fig. 4e-j; Extended Data 10). These results provide new molecular insights into the mechanism through which senescent cells regulate their persistence within tissues by modulating NK cell-mediated immunosurveillance and trigger their immune tolerance within tissues. GD3-positive cells inhibit NK cells *in vitro* and *in vivo*, while GD3-negative status is associated with NK cell cytotoxicity, both in human and murine cells. We provide the proof of the concept that GD3 expression is a senescence-associated immune checkpoint (SIC) that determines senescent cell tolerance. Moreover, we identified GD3, and SIC in general, as promising and powerful potential targets for age-associated disease therapies. Therefore, the identification and characterization of the SIC of senescence immunosurveillance opens a new avenue of research for the development of senescence-specific antibodies that may promote senescent cell clearance and thus prevent or reverse age-associated diseases.

## Methods

### Induction of senescence

Replicative senescence was induced by continuous passaging of primary human fibroblasts MRC5 at 5% O_2_ until they reached a plateau in their growth curve. Cumulative population doubling level (PDL) was calculated using the following equation: 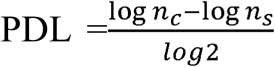 where n_c_ represents the number of cells counted after expansion and n_s_ represents the number of cells seeded. For all experiments, young cells have been used at pdl 30 and senescent cells have been defined as at least 90% SA-b-Galactosidase assay + and no more than 10% of EdU+ cells. Primary human fibroblasts MRC5 were induced to senesce by exposure to X-ray radiation at a total dose of 10 Gy at a rate of 5 Gy/min, treated with Nutlin 3a at 10 µM for 24h, CCN1 2,5 µg/mL for 6 days, Bleomycin during 24h at 50 µg/mL and Aphidicolin during 24h at 2,5 µg/mL. Oncogene-Induced senescence was induced in hMEC human mammary epithelial cells transduced with a vector expressing H-RASG12V. Cells were treated with doxycycline hyclate (Sigma-Aldrich) at 1µg/mL to activate H-RASG12V during 5 or 12 days. Mycoplasma test was performed every month, and experiments are performed only on mycoplasma negative cells.

### Animals. Matrigel Plug assays

Experiments are performed on 8- to 10-week-old female NMRI Nude mice from Charles River. 500 000 cells in 100µl of PBS or the equivalent of the supernatant of the same cells concentrated by Amicon 10KD (Milipore) in 100 µl of PBS and complemented with 400 ul of Matrigel growth factor reduced (Corning) were inoculated subcutaneously into the back under isoflurane anesthesia. Five days after, mice were sacrificed and Matrigel plugs were harvested. Infiltrating cells were collected after enzymatic dissociation by Dispase (Corning), collagenase A, and DNAse I (Roche) digestion for 30 min at 37°C. Infiltrating cells stained with directly coupled antibodies for 30 min at 4°C after saturation with Fc-Block anti-CD16/CD32 antibodies (clone 2.4G2) for 15 min on ice. After washes in 0.5 mM EDTA 2% FCS PBS, cells were analyzed using an ARIA III cytometer BD Biosciences with DIVA6 software and FlowJo 10 software. **Mouse model of ADR-induced kidney injury**. A single dose of 12 mg/kg of Adriamycin (ADR, Doxorubicin Hydrochloride, Sigma, Ref# D1515) was injected in tail vein of 3 months old BALB/c mice (Charles River). Littermates were injected similarly with saline (NaCl 0.9%). All mice were weighed twice a week in the time course of the experiments. Kidneys of the mice were collected 14 days after injection for further histological analysis. BALB/c mice were obtained from Janvier Lab (strain BALB/cJRj). **Bleomycin induced lung fibrosis**. 8- to 10-week-old pathogen-free female C57BL/6 mice (Charles River) were treated with intratracheal instillation of 50 µl of phosphate-buffered saline (PBS) or bleomycin (2.5 U.kg−1, Sigma-Amdrih). Animals were maintained in a 12:12-h light-dark cycle with food and water ad libitum. After the experiments, mouse lung tissues were excised and either included in OCT frozen, or inflated with 4% (w/v) paraformaldehyde in PBS, embedded in paraffin, sectioned, and stained with H&E and picrosirius red. The percent of tissue area that was classified as fibrosis was quantified with Image J. Alternatively, lungs were freshly dissociated with Miltenyi Lung dissociation Kit (ref 130-905-927) and GentleMacs with Heaters. Infiltrating cells stained with directly coupled antibodies for 30 min at 4°C after saturation with Fc-Block anti-CD16/CD32 antibodies (clone 2.4G2) for 15 min on ice. After washes in 0.5 mM EDTA 2% FCS PBS, cells were analyzed using an ARIA III cytometer BD Biosciences with DIVA6 software and FlowJo 10 software. All mouse experiments were conducted according to local and international institutional guidelines and were approved by either the Animal Care Committee of the IRCAN and the regional (CIEPAL Côte d’Azur Agreements NCE/2015-266#2015102215087555 and NCE/2020-675# 2020042723583497) and national (French Ministry of Research) authorities.

### Senescence-associated β-galactosidase staining on cultured cells

Cells were fixed and stained using the Senescence-Associated β-Galactosidase Staining Kit (Abcam ab65351), following the manufacturer’s instructions. After staining, cells were incubated for 12-14h at 37 °C, then visualized by phase-contrast microscopy. The percentage of Senescence-Associated β-Galactosidase-positives cells is calculated by determining the ratio of Senescence-Associated β-Galactosidase-positives cells (blue staining) among all cells counted. A minimum of 150 cells is counted for each condition at each experiment using the proliferative MRC5 (PDL30) as a negative control.

### EdU proliferation assay on cultured cells

To measure cell proliferation, an EdU (5-ethynyl-2′-deoxyuridine) proliferation assay was performed as recommended by Thermo Fisher Scientific. MRC5 and hMEC cells were plated in 24-well plates at a density of 5×10^4^ cells/well for 12 hours. Then incubated in 50% conditioned medium and 50% serum-free DMEM containing 10μmol/L EdU (Invitrogen/ Click-IT EdU Alexa Fluor 647 Imaging Kit) for 12hrs. Cells were fixed, then underwent DNA staining, according to the manufacturer’s instructions to detect the number of cycling cells during the EdU treatment. The cells were imaged using confocal fluorescence microscopy (Zeiss LSM880) and the number of proliferating cells was averaged to calculate the percentage of proliferative cells. The percentage of proliferative cells is determined by calculating the ratio of the number of proliferating cells (EdU positive/red nucleus) to the total number of cells counted (DAPI+). A minimum of 100 cells is counted for each condition at each experiment using the proliferative MRC5 (PDL30) as a positive control.

### Extraction and purification of glycolipids

Cells (2×10^7^) were lyophilized and extracted three times with CHCl_3_/CH_3_OH (2:1, *v/v*) and once by CHCl_3_/CH_3_OH (1:2, *v/v*) using intermediary centrifugations at 2500 g for 20 min. Combined supernatants were dried under a nitrogen stream, subjected to mild saponification in 0.1 M NaOH in CHCl_3_/CH_3_OH (1:1, *v/v*) at 37°C for 2 h and evaporated to dryness. Samples were reconstituted in CH_3_OH/0.1% TFA in water (1:1, *v/v*) and applied to a reverse phase C_18_ cartridge (Waters, Milford, MA, USA) equilibrated in the same solvent. After washing with CH_3_OH/0.1% TFA in water (1:1, *v/v*), GSL were eluted by CH_3_OH, CHCl_3_/CH_3_OH (1:1, *v/v*) and CHCl_3_/CH_3_OH (2:1, *v/v*). The elution fraction was dried under nitrogen stream prior to structural analysis.

### Sequential release and purification of *N*- and *O-*glycans

Cells were resuspended in 6 M guanidinium chloride and 5 mM EDTA in 0.1 M Tris/HCl, pH 8.4, and agitated for 4 h at 4 °C. Then, dithiothreitol were added to a final concentration of 20 mM and incubated during 5 h at 37°C, followed by addition of 50 mM iodoacetamide overnight in the dark at room temperature. The reduced/alkyled glycoproteins were dialyzed against water at 4°C for 72 h and lyophilized. Samples were incubated with trypsin TPCK (Sigma-Aldrich) at a 20:1 ratio *(w/w)* in 50 mM NH_4_HCO_3_, pH 8.5, for 24 h at 37°C. The digestion was stopped by incubation at 100°C for 5 min, followed by C_18_ Sep-Pak chromatography (Waters Corp., Guyancourt, France). C_18_ Sep-Pak was equilibrated in 5% aqueous acetic acid and washed in the same solvent. Sample was loaded on the C_18_ Sep-Pak and the bound peptides were eluted with 20%, 40% and 60% *(v/v)* propanol in 5% aqueous acetic acid, pooled and lyophilized. *N-*Glycans were released by 10 U *N-*glycosidase F (Roche) digestion in 50 mM NH_4_HCO_3_ buffer pH 8.4, overnight at 37°C. *N-*glycans and *O-*glycopeptides were separated by C_18_ Sep-Pak, following the same protocol described above. Propanol fractions, containing *O-*glycopeptides, were pooled and freeze-dried. To liberate *O-*glycans, peptides were submitted to reductive elimination in 1 M NaBH^4^ and 0.1 M NaOH at 37°C for 72 h. The reaction was stopped by the addition of Dowex 50×8 cation-exchange resin (25−50 mesh, H^+^ form) until pH 6.5 was reached. After evaporation to dryness, boric acid was distilled in the presence of methanol. Total material was then submitted to cation-exchange chromatography on a Dowex 50×2 column (200−400 mesh, H^+^ form) to remove residual peptides.

### Mass Spectrometry

Glycans and glycolipids were permethylated according to the method of Ciucanu and Kerek^21^ prior to mass spectrometry analysis. Briefly, samples were incubated with DMSO/NaOH/ICH_3_ during 2 h under sonication. The reaction was stopped with water and the permethylated glycans were extracted in CHCl_3_ and washed at least seven times with water. Permethylated glycans were solubilized in CHCl_3_ and mixed with 2,5-dihydroxybenzoic acid matrix solution (10 mg/mL dissolved in CHCl_3_/CH_3_OH (1:1, v/v) and spotted on MALDI plate. MALDI-TOF mass spectra were acquired on Voyager Elite DE-STR mass spectrometer (Perspective Biosystems, Framingham, MA, USA) and MALDITOF/TOF analyzed on 4800 Proteomics Analyzer mass spectrometer (Applied Biosystems, Framingham, MA, USA) in reflectron positive mode by delayed extraction using an acceleration mode of 20 kV, a pulse delay of 200 ns and grid voltage of 66%. For each spectrum, 5000 laser shots were performed and accumulated.

### Immunofluorescence

To analyze the presence of the ganglioside GD3 at the cell surface of MRC5, we seed 5.10^4^ cells in the 24 wells plate. Cells were fixed during 10 minutes at room temperature with the 1X PBS solution containing 4% Formaldehyde (Sigma). For the GD3 staining, we use a primary antibody anti-GD3 R24 (Abcam) at 1:1000, overnight at 4°C. A secondary antibody against mouse whole IgG in FITC is used at 1:3000 during 1 hour at room temperature (Jackson ImmunoResearch). The autofluorescence of senescent cells was reduced by the utilization of Autofluorescence Eliminator Staining (Merck Millipore) staining following the manufacturer’s instructions. The analysis was performed using fluorescence microscopy and used the same time of laser exposure between all the conditions.

### Western blotting

Cells were harvested by trypsinization, washed in PBS and lysed with radio-immunoprecipitation assay (RIPA) buffer (Sigma-Aldrich) supplemented with protease and phosphatase inhibitors (Sigma-Aldrich) for 30 min on ice. Protein concentrations were determined using the BCA Protein Assay Kit (Interchim). Cell lysates (40 μg of the total protein) were diluted in SDS sample buffer with reducing agent (NuPage, Life Technologies) and boiled for 5 min at 95 °C. Cell lysates were separated by protein electrophoresis at 150 V for 1h using 4-20% Mini-Protean TGX pre-cast gels (BioRad) and transferred by semi-dry technic onto Amersham Hybond-P PVDF membranes (GE Healthcare). After blocking, membranes were probed with primary antibodies overnight at 4 °C, washed and incubated with HRP-conjugated secondary antibodies (Vector, 1:20000) for 1h at room temperature. Antibodies were detected using the ECL detection kit (GE Healthcare). Prior to re-probing with different antibodies, membranes were stripped at 4°C in agitation using Antibody stripping buffer 1X for 10 minutes (Gene Bio-Application). Protein bands were quantified using ImageJ software. The integrated density of each band was measured using the gel analysis function of ImageJ, normalized to alpha-tubulin. Primary antibodies were mouse monoclonal anti-ganglioside GD3 R24 (Abcam), mouse monoclonal alpha-tubulin (Sigma).

### Quantitative real-time PCR

Total RNA isolation from cells was performed using Trizol (Sigma-Aldrich). Reverse transcription (RT) was performed with the High-Capacity RNA to DNA kit (Applied Biosystems). Quantitative real-time PCR was performed on a Step-One Plus real-time system (Applied Biosystems) according to the manufacturer’s protocol. qPCRs were made on cDNAs obtained using Roche’s Fast Start Universal SYBR Green. Data were analyzed according to the Pfaffl method after calculation of primer efficiency. RPL0 was used as an endogenous control. All reactions were performed in triplicate, and at least 3 independent experiments were performed to generate each data set. The primers were as follows: hST8SIA1 sense 5’-GGGTGAGGCAAGTTGAAAGG-3’ – antisense 3’-AGGTCCTCAGCGAATTTCCA-5’; RPL0 sense 5’ -AACTCTGCATTCTCGCTTCCT-3’ – antisense 3’-ACTCGTTTGTACCCGTTGATG-5’

### GD3 dosage by ELISA

Serum from young (3 months-old) or old (24 months -old) mice or supernatant from young MRC5 until replicative senescent MRC5 were collected. The amount of free GD3 in sera or supernatants were assessed by ELISA against GD3 following the recommendation of the manufacturer (CUSABIOTECH ref CSB-EQ027866HU).

### Coculture experiment

Splenocytes are extract from C57Bl/6j naïve mice previously stimulated *in vivo* for 14h by an intraperitoneal injection of 150 µg of Poly I:C (Invivogen). The MRC5 are plated at 5×10^4^ cells/well in 48 wells plate the day before the experiment for the good adherence of the cells. All the coculture were performed with four different mice at each time. Then, the killing capacity of NK cells was tested when they coculture with senescent / young or modulated for ST8SIA1 expression MRC5.NK cells are added to the culture for 4 h in presence of monensin and brefeldin (BD Biosciences) at the effector/target ratio of 1:1. Degranulation activity of the NK cells is then measured by FACS by the anti-CD107a (FITC BD Biosciences) and IFN-γ staining (PE BD Biosciences).

### *ex vivo* rechallenge experiment

Primary cells from the spleen (crushing on cell strainer 70µm) or from the lungs (dissociation with Miltenyi Lung dissociation Kit ref 130-905-927 and GentleMacs with Heaters) were extracted from PBS or Bleomycin instillated mice. Bulk of primary cells (either from the spleen or the lungs) were then rechallenge in vitro with YAK-1 cells for 4 hours in presence of monensin and brefeldin (BD Bioscience) at the Effector/Target ratio of 5: 1. Degranulation activity of the NK cells is then measured by FACS by the anti-CD107a and IFN-γ staining.

### Real-time NK cell cytotoxic assay

A real-time cytotoxic assay was performed as previously described^22,23^. Briefly, target cells were labelled with 0.5µM Calcein-AM (Molecular Probes) for 15 min at room temperature. Proliferative or repmicative senescent MRC5 cells were additionally treated with 100 µM Indomethacin (Sigma Aldrich) to block multidrug-resistance transporters that expulse calcein. The inhibitor was maintained in the medium during the assay. Human primary NK cells purified from PBMC from healthy donors after FACS sorting. Calcein-labelled targets were incubated with human NK cells for 4 h at 37°C, 5% CO2 and real-time monitoring of NK cell killing was performed on a Cytation(tm) 5 (Biotek). Cell images were processed using GEN5 software (Biotek). The percentage of lysis from triplicates was calculated as follow: % lysis = {1-[(experimental well at t/experimental well at t0)/(control well at t/control well at t0)]} × 100.

### *In-vivo* lung imaging by computed tomography (µCT)

High-resolution CT scan were performed using a dedicated system (eXplore speCZT CT120, GE Healthcare). Mice were gas anesthetized (air and 1-2% isoflurane) in an air-warmed imaging chamber (Minerve) to maintain body temperature during the scanning time. Micro-CT image acquisition consisted of 400 projections collected in one full rotation of the gantry in approximately 5 min in a single bed focused on the lungs, with a 450 mA/80kV X-ray tube. 2-D and 3-D images were obtained and analyzed using the software program MicroView (GE Healthcare).

### Senescence-associated β-galactosidase staining and DNA Damage analysis at telomeres (Telomere Induced Foci) by ImagestreamX analysis

Cells were resuspended (either cells in culture or primary cells after tissues digestion) in phosphate□buffered saline (PBS), then were fixed with 4% paraformaldehyde (PFA) for 15 min at room temperature and stained
using the Senescence-Associated β-Galactosidase Staining Kit (Abcam ab65351), following the manufacturer’s instructions. After staining, cells were incubated for 12-14h at 37 °C sealed and protected from light. After washes, cells were analyzed by ImageStreamX or permeabilized with PBS-TritonX100 (0.5%) and incubated 10 minutes at RT, then at 87°C for 10 minutes while vortexing, and overnight at room temperature protected from light with (70% formamide, 1% blocking reagent, 10mM Tris ph7.2, 4nM PNA probe-FITC). Cells were stained with specific antibody for 53BP1 (1/300) (Novus Biological) then incubated with Goat anti Rabbit Alexa 647 (1/900) (Jackson ImmunoResearch). Nucleus were stained with Hoeuchst (1/2000) then cells were washed twice and analyzed by ImageStreamX.

### Histology & Immunohistochemistry

Antigen retrieval was performed on 5μm paraffin sections using Vector unmasking reagent (Vector Laboratories, Ref# H3300). Tissue sections were blocked (MOM kit, Vector Laboratories, Ref# BMK-2202), and incubated with mouse monoclonal anti-GD3 antibody (Abcam ab11779, 1:40) for overnight at 4°C. Primary antibody was detected using a biotinylated anti-mouse IgG (MOM kit, Vector Laboratories, Ref# BMK-2202) followed by streptavidin-Cy3 (Jackson ImmunoResearch, Ref# 016-160-084). For fibrosis analysis, slides were stained with picro-sirius red solution for 1h and washed with acetic acid solution and absolute alcohol before imaging in white light or polarized light.

Image analysis using ImageJ software. Stained tissue sections were sequentially scanned using an HD Zeiss microscope allowing imaging of the entire section. For signal analysis, all glomeruli (about 120) were manually demarcated within each kidney section. Quantification of the signal within glomeruli or within the remaining area of the kidney was performed using Image J software.

### Antibodies

| Specificity | Company | Clone | Species | Isotype | Fluorochrome | Reference |
| --- | --- | --- | --- | --- | --- | --- |
| **anti Ly6G (Gr1)** | eBioscience | RB6-8C5 | rat | IgG2b, k | PE | 12-5931-82 |
| **anti CD107a** | BD Biosciences | 1D4B | Rat | IgG2a/k | FITC | 553793 |
| **anti CD11b** | BD Biosciences | M1/70 | Rat | IgG2b | APC-H7 | 550993 |
| **Anti CD11c** | BD Biosciences | HL3 | Hamster | IgG1 | FITC | 557400 |
| **anti CD19** | BD Biosciences | 1D3 | rat | IgG2a, κ | FITC | 553785 |
| **Anti NKp46** | BD Biosciences | 29A1.4 | Rat | IgG2a | PE | 560757 |
| **Anti CD45** | BD Biosciences | 30-F11 | Rat | IgG2b | PerCP | 557235 |
| **Anti CD45** | BD Biosciences | 30-F11 | Rat | IgG2b | A700 | 560510 |
| **Anti CD69** | BD Biosciences | H1.2F3 | Hamster | IgG1/K | PE-Cy7 | 552879 |
| **anti CD8a** | BD Biosciences | 53-6.7 | rat | IgG2a,K | BV650 | 563152 |
| **anti IFN-γ** | BD Biosciences | 4S.B3 | Mouse | IgG1/K | PE | 554552 |
| **anti IFN-γ** | BD Biosciences | XMG1.2 | Rat | IgG1/K | PE | 554412 |
| **anti NK-1.1** | BD Biosciences | PK136 | Mouse | IgG2a/k | APC | 550627 |
| **anti NK-1.1** | Biolegend | PK136 | Mouse | IgG2a, κ | APC | 108710 |
| **Anti NKp46** | BD Biosciences | 29A1.4 | Rat | IgG2a | Alexa 647 | 560755 |
| **anti-CD107a** | BD Biosciences | 1D4B | rat SD | IgG2a, κ | V450 | 560648 |
| **Anti-CD11b** | Biolegend | M1/70 | Rat | IgG2b, κ | BV605 | 101237 |
| **anti-CD11c** | Biolegend | N418 | Hamster | IgG | PE/dazzle | 117347 |
| **anti-CD19** | Biolegend | 6D5 | rat | IgG2a, κ | BV510 | 115545 |
| **Anti-CD19** | Biolegend | 6D5 | Rat | IgG2a, κ | BV785 | 115543 |
| **Anti-CD25** | eBioscience | PC 61.5 | Hamster | IgG1 λ | PE-Cy7 | 25-0251-82 |
| **anti-NKp46** | BD HORIZON | 29A1.4 | Rat | IgG2a, κ | BV510 | 563455 |
| **anti-CD3e** | BD Biosciences | 145-2C11 | Hamster | IgG1, κ | FITC | 553062 |
| **Anti-CD3e** | Biolegend | 145-C11 | Hamster | IgG | PerCP | 100302 |
| **anti-CD4** | BD Biosciences | GK1.5 | rat | IgG2b, κ | PE | 553730 |
| **anti-CD4** | LifeTech | monoclonal | rat | IgG2a | PE-AF700 | MCD0424 |
| **anti-CD8a** | BD Biosciences | 53-6.7 | Rat LOU | IgG2a, κ | PerCP-Cy5.5 | 551162 |
| **Anti-F4/80** | Biolegend | BM8 | Rat | IgG2a, κ | PerCP-Cy5.5 | 123128 |
| **Anti-F4/80** | Biolegend | BM8 | Rat | IgG2a, κ | BV510 | 123135 |
| **Anti-Ly6C** | eBiosciences | HK1.4 | rat | IgG2c, κ | APC-eFluor780 | 47-5932-82 |
| **Anti-Gr-1** | Biolegend | RB6-8C5 | rat | IgG2b, κ | PE | 108408 |
| **Anti-Gr-1** | BD Biosciences | RB6-8C5 | Rat | IgG2b, κ | PE | 553128 |
| **Anti-Gr-1** | Biolegend | RB6-8C5 | Rat | IgG2b, κ | BV421 | 108434 |
| **Anti-Ly6G** | BD Biosciences | 1A8 | Rat LEW | IgG2a, κ | PE-Cy7 | 560601 |
| **Anti-Ly6G** | Biolegend | 1A8 | Rat | IgG2a, κ | BV421 | 127627 |
| **Anti-Ly6G** | BD Biosciences | 1A8 | Rat LW | IgG2a, κ | BV711 | 563979 |
| **anti-GD3** | Abcam | R24 | mouse | IgG3 | uncoupled | ab11779 |
| **Anti-GD3** | Biotem | R24 Hybridoma ATCC HB-8445 | mouse | IgG3 | Uncoupled endotoxin free for *in vivo* experiment | Hybridoma ATCC HB-8445 |
| **Anti-53BP1** | Novus Biological | Rabbit polyclonal | Rabbit | IgG | uncoupled | NB100-305 |
| **Recombinant Human Siglec-7 Fc Chimera Protein** | R&D Systems | Human recombinant |  |  | uncoupled | 1138-SL-050 |

### GTEX human gene expression analysis

The Genotype-Tissue Expression (GTEx) project is an ongoing effort to build a comprehensive public resource to study tissue-specific gene expression and regulation. Samples were collected from 54 non-diseased tissue sites across nearly 1000 individuals. We used the raw open access files from GTEX datasets to extract genes expression and various metainformations such as age, sex or organs origin from the human donor’s cohort. We generated a home-made Python script (freely available on demand) that crossed GTEX RNA-seq data with related GTEX annotations files. The script is usable under Windows, Mac or Linux. The output is saved in a single Excel file containing the information for the selected gene. Data were then plotted and statistically analyzed using Prism GraphPad version 8.

### Statistical analysis

Reasonable sample size was chosen to ensure adequate reproducibility of results and was based on our previous studies. Mouse experiments are performed on n = 4 individuals as indicated in Fig legends. Statistical tests were all performed with GraphPad Prism 8 software including the Student’s t-test, the Mann– Whitney test, the log-rank test, Pearson correlation, and two-way ANOVA tests.

## Acknowledgment

We thank the IRCAN core facilities (animal facility, CytoMed, GenoMed and), which are supported by le FEDER, Ministère de l’Enseignement Supérieur, Région Provence Alpes-Côte d’Azur, Conseil Départemental 06, ITMO Cancer Aviesan (plan cancer), Cancéropole PACA, the Association pour la Recherche sur le Cancer (ARC), the Infrastructures en Biologie Santé et Agronomie (IBiSA), l’Inserm and PAGes core facili<u>ty (ht</u>tp:/<u>/plateforme-</u>pages.univ-lille1.fr) and UMS 2014 - US 41 - Plateformes Lilloises en Biologie & Santé for providing the scientific and technical environment conducive to achieving this work.

## Author’s contributions

J.C.V. conceived the study with the support of EG; C.I, E.G. and J.C.V. designed experiments; C.I., L.S., L.C., T.H., L.M., M.-C.M., S.K., O.C., and J.C.V. conducted experiments; C.D. Y.G., F.A. performed mass spectrometry analysis; J.G and T.P performed in vivo µCT bioimaging; C.I., Y.G., F.A., M.-C.M., L.S., C.F., E.G., and J.C.V. analyzed data; E.G and J.CV secured funding. J.C.V. and E.G. wrote the manuscript. All the authors read and approved the paper.

## Additional information

Correspondence and requests for materials should be addressed to E.G. and J.C.V.

## Extended data figure legends

**Extended Data 1: Characterization of MRC5 replicative senescent cells. a**, Growth curve of the MRC5 cells until replicative senescence. pdl. Population doubling level. **b**, SA-β−Galactosidase and EdU incorporation assay on pdl30 or replicative senescent MRC5 (left panel) and quantification of the percentage of SA-βGalactosidase + and EdU – cells (right panels). **c-e**, Analysis of the SA-β−Galactosidase assay, DNA damage level (53BP1 staining) (**c**,**d**) or DNA damage level(53BP1 staining) specifically at telomeres (telomere staining (PNA-Telo) and colocalization (TIF for Telomere Induced Foci)) (**d**,**e**) on pdl30 or replicative senescent MRC5 using ImagestreamX and quantification of the number of total 53BP1 foci or 53BP1 foci at telomere (TIF). Due to the SA-b-Galactosidase staining, cells are darker, and the Mean pixel intensity is decreasing (**e**). Data are representative of 8 different batches of replicative senescent cells corresponding to the batches used in Fig.1 to Fig. 3.

**Extended Data 2: Senescent cells but not their SASP affect immune infiltration and NK cell activation. a**, Quantification of immune cells infiltrating the Matrigel plugs assay in presence of pdl30 or replicative senescent cells (corresponding to Fib. 1b). **b**, representative FACS dot plot of the total immune (CD45+), NK cell (NKp46+ cells) or myeloid cells (CD11b+ GR1+) infiltration. **c**, representative scheme of the Matrigel plug assay with only conditioned media of MRC5 cells. **d-e**, Quantification of immune cells infiltrating the Matrigel plugs assay in presence of pdl30 or replicative senescent cells supernatant. **f**, positive (YAC-1 cells, a strong NK cell target) and negative controls of degranulation (CD107a+ NK cells) or IFN-γ production in *in vitro* co-culture experiments. **g**, analysis of IFN-γ production in *in vitro* co-culture experiments with senescent cells. All experiments are performed with n = 4 mice per group (**a-e**); *p < 0.05, **p < 0.01, and ***p < 0.001; Mann–Whitney *U* test or data represent the mean of n = 3 independent experiments; *p < 0.05, **p < 0.01, and ***p < 0.001; Mann–Whitney test (**f-e**).

**Extended Data 3: Mass spectrometry analysis of human replicative senescent cells for O-glycans. a**, scheme representing the analysis strategy. **b-c**, representative data of the O-glycan composition of pdl30 (young) MRC5 (**b**) or replicative senescent MRC5 (**c**).

**Extended Data 4: Mass spectrometry analysis of human replicative senescent cells for permethylated N-glycans. a-b**, representative data of the N-glycan composition of pdl30 (young) MRC5 (**a**) or replicative senescent MRC5 (**b**).

**Extended Data 5: Replicative and Stress induced senescent cells reshuffled their ganglioside composition. a**, quantification of GD3 expression analysis by cytometry (corresponding to Fig. 2b). **b**, representative scheme of ganglioside biosynthesis pathway. **c**, qPCR analysis of ganglioside biosynthesis enzymes in replicative senescent cells (representative data of 3 independent experiments). **d**, growth curve of MRC5 and kinetic of *ST8SIA1* expression by qPCR during replicative senescence. **e**, GD3 dosage by ELISA in supernatants of proliferating or replicative senescent cells (2 independent experiments). **f**,**g**, qPCR analysis of *ST8SIA1* expression in replicative and stress induced senescence (**f**) and of ST8SIA1 protein level by western-blot (**g**) (representative data of 3 independent experiments).

**Extended Data 6: Human and mouse Oncogene Induced Senescent (OIS) cells repress *ST8SIA1* and do not express the GD3 ganglioside. a-b**, Morphology analysis and SA-β−Galactosidase assay and quantification for HMEC cells overexpressing Hras in inducible manner. **c-d**, analysis by qPCR of *ST8SIA1* expression in HMEC overexpressing Hras senescent cells or Bleomycin induced HMEC senescent cells (**c**) or in replicative senescent or RasV12 induced senescent WI38 cells (**d**). **e**, SA-β−Galactosidase assay and GD3 immunofluorescence staining on paraffin embedded lung section from control or 2 months-induced KrasG12D overexpression mice. **f**, Analysis of GD3 expression in lungs of KrasG12D overexpression mice overtime.

**Extended Data 7: The expression of a sialylated GD3 is strictly required for inhibition of NK cell degranulation by senescent cells. a**, Cytometry analysis of the binding of soluble recombinant Siglec-7-Fc proteins by pdl30 or replicative senescent MRC5 cells. **b**, analysis by qPCR of *ST8SIA1* expression in replicative senescent after lentiviral infection for shRNA targeting *ST8SIA1*. **c**, Quantification of the percentage of SA-β-Galactosidase positive and EdU negative cells after *ST8SIA1* knock-down. **d**, Flow cytometry analysis of GD3 expression in replicative senescent after lentiviral infection for shRNA targeting *ST8SIA1*. **e-f**, cytometry analysis of GD3 expression (**e**) and qPCR expression of *ST8SIA1* (**f**) by pdl30 MRC5 overexpressing or not *ST8SIA1*. **g**, Quantification of the percentage of SA-β-Galactosidase positive and EdU negative cells after ST8SIA1 overexpression. **h**, IFN-γ production by NK cells in *in vitro* co-culture experiments with pdl30 or replicative senescent MRC5 cells treated or not with neuraminidase. **i**, degranulation (CD107a+; left panel) or IFN-γ production (right panel) by NK cells in *in vitro* co-culture experiments using pdl30 MRC5 overexpressing ST8SIA1. Data represent the mean of n = 3 independent experiments; *p < 0.05, **p < 0.01, and ***p < 0.001; Mann–Whitney *U* test.

**Extended Data 8: mAb against GD3 restore efficient NK cell degranulation and killing of senescent cells. a**, Dose response effect of GD3 mAb on NK cell degranulation against replicative senescent or pdl30 MRC5 (data corresponding to Fig. 3e). **b**, IFN-γ production in *in vitro* co-culture experiments using anti-GD3 monoclonal antibody (corresponding to Fig. 3e). **c-d**, degranulation (**c**, CD107a+ NK cells) or (**d**) IFN-γ production in *in vitro* co-culture experiments using isotypic mouse IgG3 antibody as control as same concentration than anti-GD3 mab. **e**, *in vitro* killing assay of pdl30 MRC5 or replicative senescent MRC5 by human NK cells in presence or not of anti-GD3 mAb; performed in same time than experiment in Figure 1f, same controls. Data represent the mean of n = 3 independent experiments; *p < 0.05, **p < 0.01, and ***p < 0.001; Mann–Whitney *U* test.

**Extended Data 9: The accumulation and long-term persistence of GD3 positive senescent cells within experimental Bleomycin-induced lung fibrosis inhibits NK cell functionality. a-b**, H&E or Sirius red staining (in white or polarized light) in bleomycin induced lung fibrosis sections (**a**) and quantification of collagen deposit (Sirius red staining; **b**). **c**, SA-β-Galactosidase assay and GD3 expression analysis (**b**) corresponding to fibrotic lungs used in Fig. 4a-d. **d-e**, histology analysis, quantification of collagen deposit (Sirius red staining in white or polarized light) and GD3 expression in fibrotic lungs over time (**d**) and their quantification (**e**). **f**, analysis of the immune infiltration by flow cytometry of control or fibrotic lungs used in Fig. 4a-d. **g**, *ex vivo* co-culture experiment using splenocytes from control or bleomycin instillated mice against the oncogenic NK cell target YAC-1 cells. All experiments are performed with n = 8 mice per group; *p < 0.05, **p < 0.01, and ***p < 0.001; Mann–Whitney *U* test.

**Extended Data 10: anti GD3 targeting *in vivo* restore NK cell functionality locally *in vivo* and *ex vivo*. a-b**, Analysis of the GD3 expression intensity by ImagestreamX (**a**) and of the percentage of GD3+ senescent cells (**b**). Data corresponding to Fig. 4h. **c**, analysis of the immune infiltration in the lungs by flow cytometry of isotypic control or anti GD3 mAb treated fibrotic mice used in Fig. 4e-j. **d**, evaluation by flow cytometry of the percentage of CD69+ activated intrapulmonary NK cells at d27; data corresponding to Fig. 4i. e, determination of the quantity of NK cells within the lung of fibrotic mice depending on the treatment. **f**, Determination of the intrapulmonary NK cell functionality *ex vivo* from treated or control mice against YAC-1 cells after 4 hours of rechallenge. **g**, determination of the quantity of NK cells within the spleen of fibrotic mice depending on the treatment. **h**, Determination of the NK cell functionality *ex vivo* from the spleen of treated or control mice against YAC-1 cells after 4 hours of rechallenge. All experiments are performed with n = 8 mice per group; *p < 0.05, **p < 0.01, and ***p < 0.001; Mann–Whitney *U* test.

**Extended Data 11: GD3 expression by senescent cells in experimental (Adriamycin treated mice) kidney fibrosis. a**, SA-β-Galactosidase assay and GD3 expression analysis in Adriamycin-induced kidney fibrosis. **b**, GD3 expression in kidney glomeruli from Adriamycin (ADR)-treated or control (NaCl) mice. **c**, quantification of the percentage of kidney area (left panel) or the glomerular area (right panel) covered by GD3 + signal in Adriamycin (ADR) treated or control (NaCl) kidney. All experiments are performed with n = 4 mice per group or at least 461 glomeruli; *p < 0.05, **p < 0.01, and ***p < 0.001; Mann– Whitney *U* test.

**Extended Data 12: GD3 is not secreted in the serum of old mice but is strongly increased by senescent cells found in natural Age-associated kidney fibrosis. a**, GD3 dosage by ELISA in serum of 3 months or 24 months-old mice (right panel, 8 to 9 mice per group). **b**, GD3 expression in kidney glomeruli from 6-months or 20-months old mice. **c, Q**uantification of the percentage of kidney area (left panel) or the glomerular area (right panel) covered by GD3 + signal in 6-months or 20-months old kidney. All experiments are performed with n = 4 mice per group or at least 461 glomeruli; *p < 0.05, **p < 0.01, and ***p < 0.001; Mann– Whitney *U* test.

**Extended Data 13: *ST8SIA1* expression is increased during human lung aging and correlates with the expression of senescence markers upon aging. a-b**, analysis of gene expression by RNAseq of normal human lung samples from different ages. Data are extracted from GTEX consortium and the relative gene expression (transcript per million or TPM) is represented in function of the group of age (**a**) or in box plot to compare TPM between young (20 to 29 years old) to elderlies (60 to 80 years-old) (**b**). *p < 0.05, **p < 0.01, and ***p < 0.001; Mann–Whitney *U* test. **c-d**, gene expression correlation (in TPM) between ST8SIA1 expression and p16 (**c**) or p21 gene expression (**d)**. Data represent the Pearson uncentred correlation.

