## Supplementary figures and images for "A ganglioside-based senescence-associated immune checkpoint"

### Extended Data 1

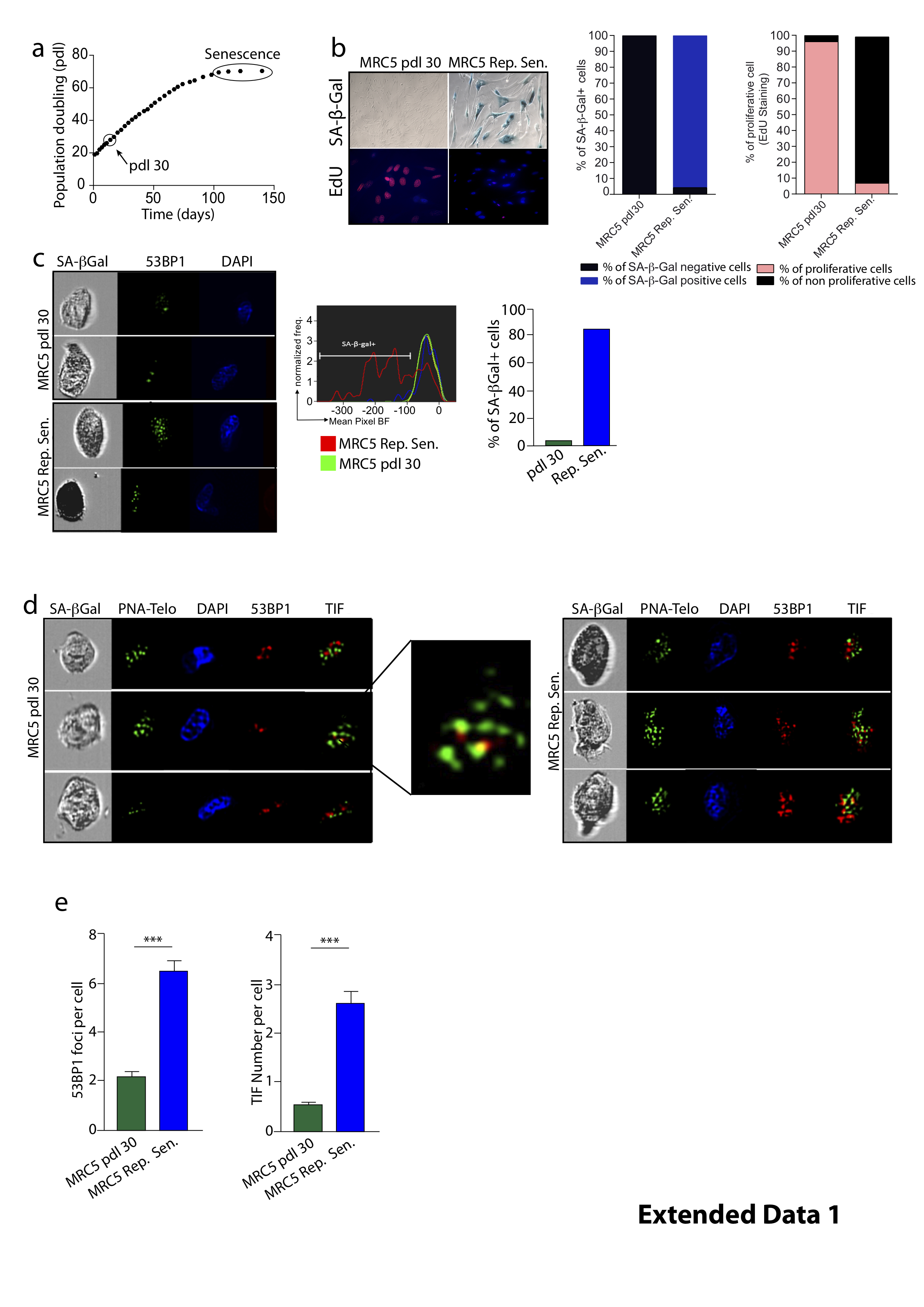

### Extended Data 2

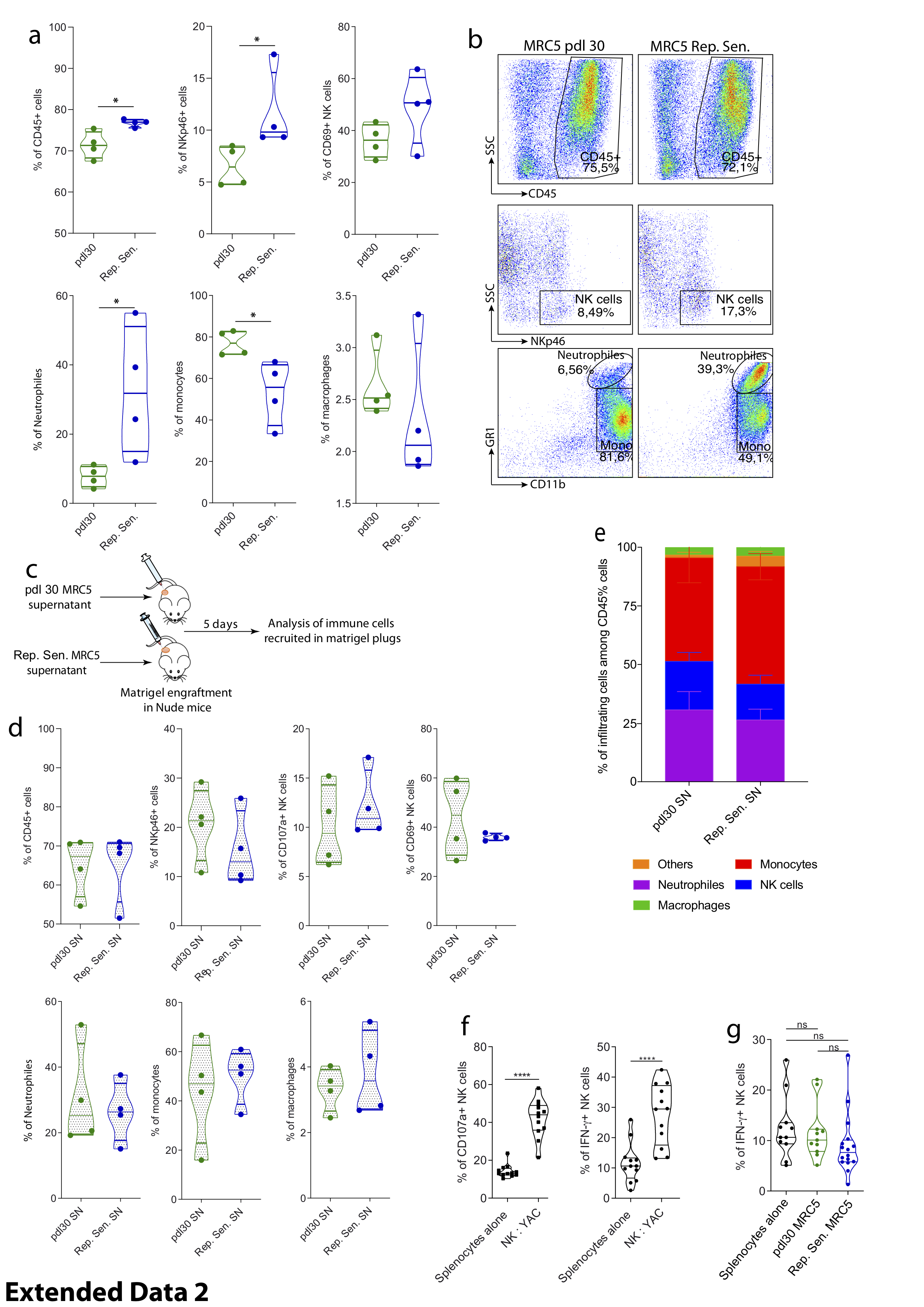

### Extended Data 3

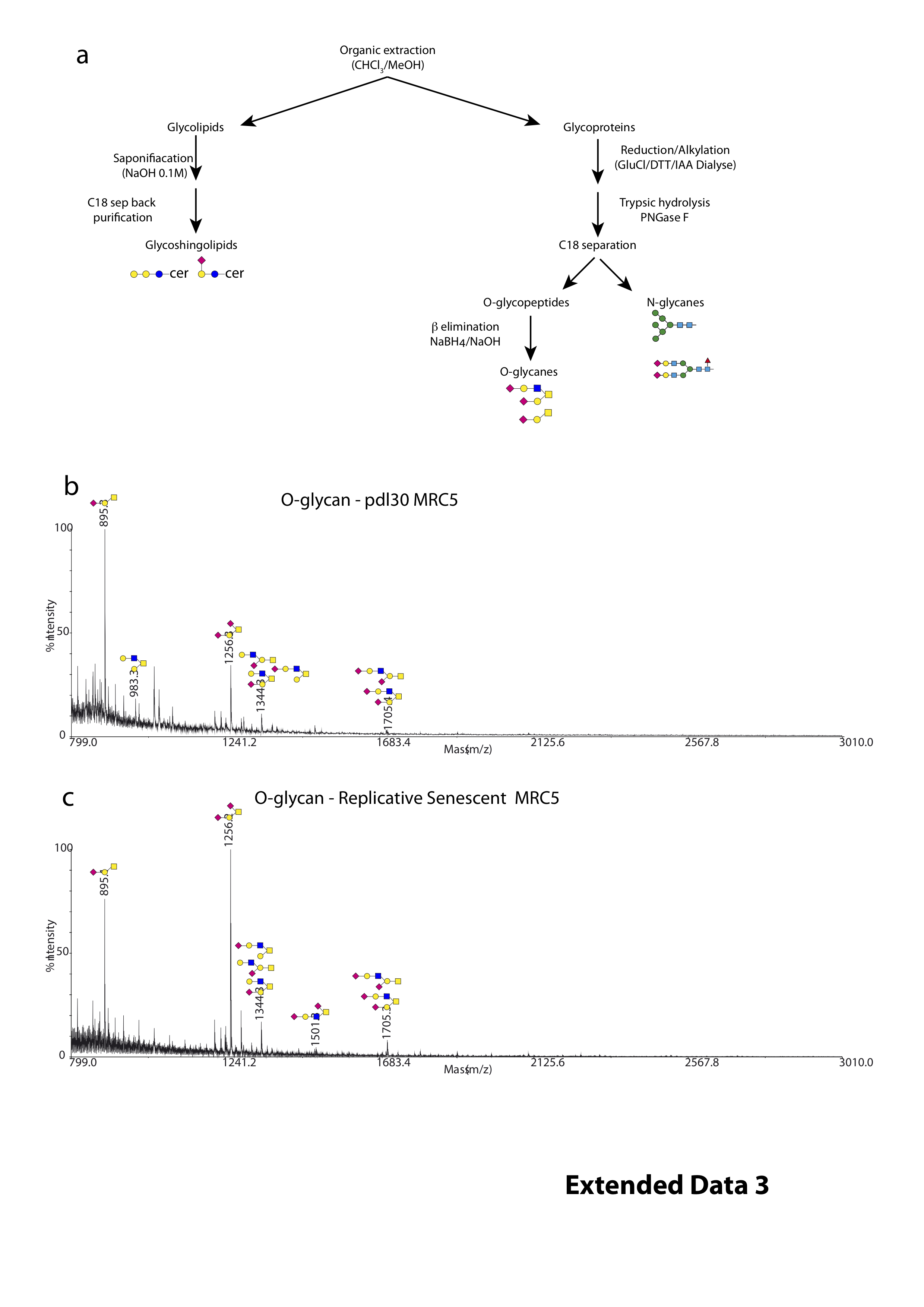

### Extended Data 4

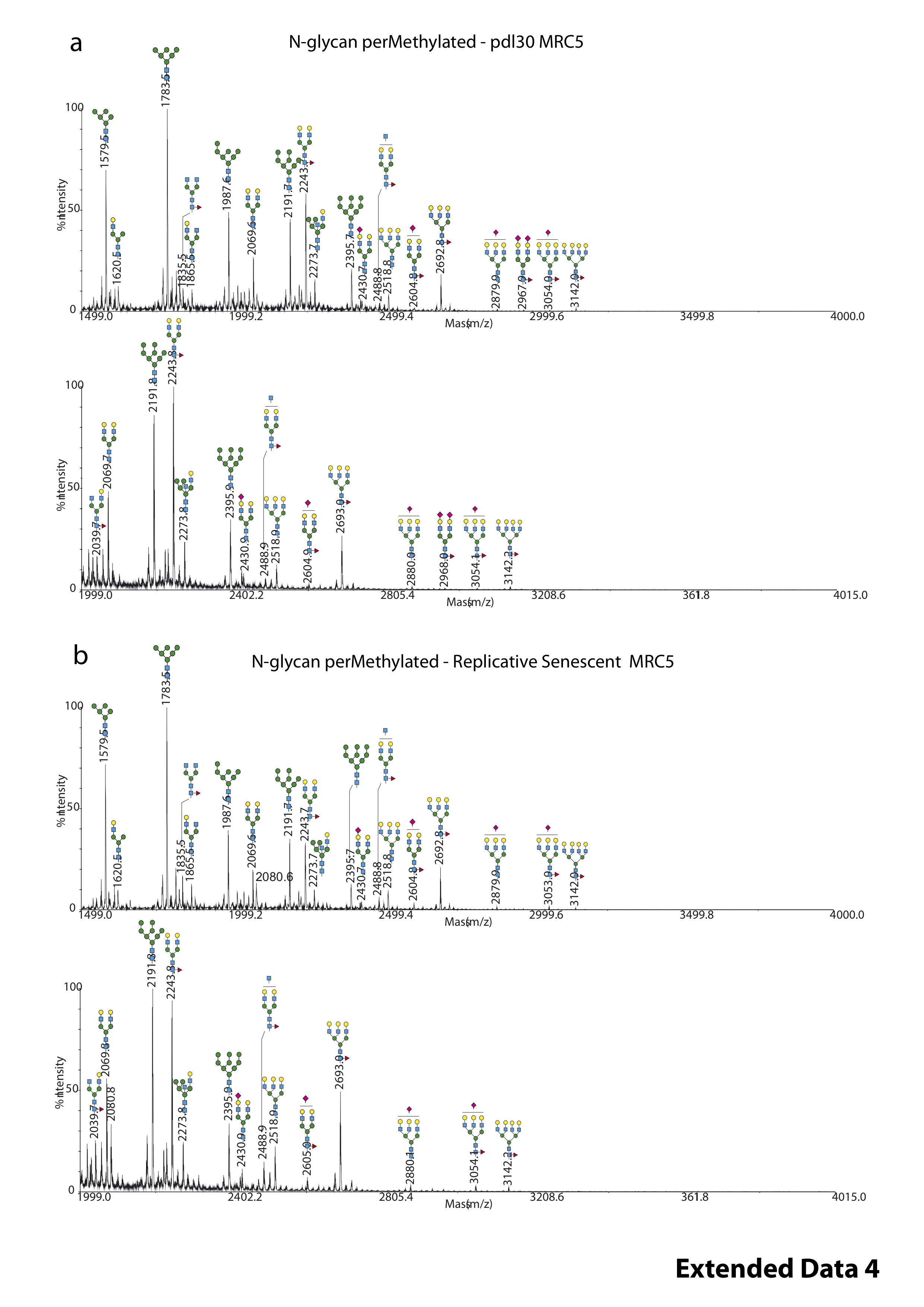

### Extended Data 5

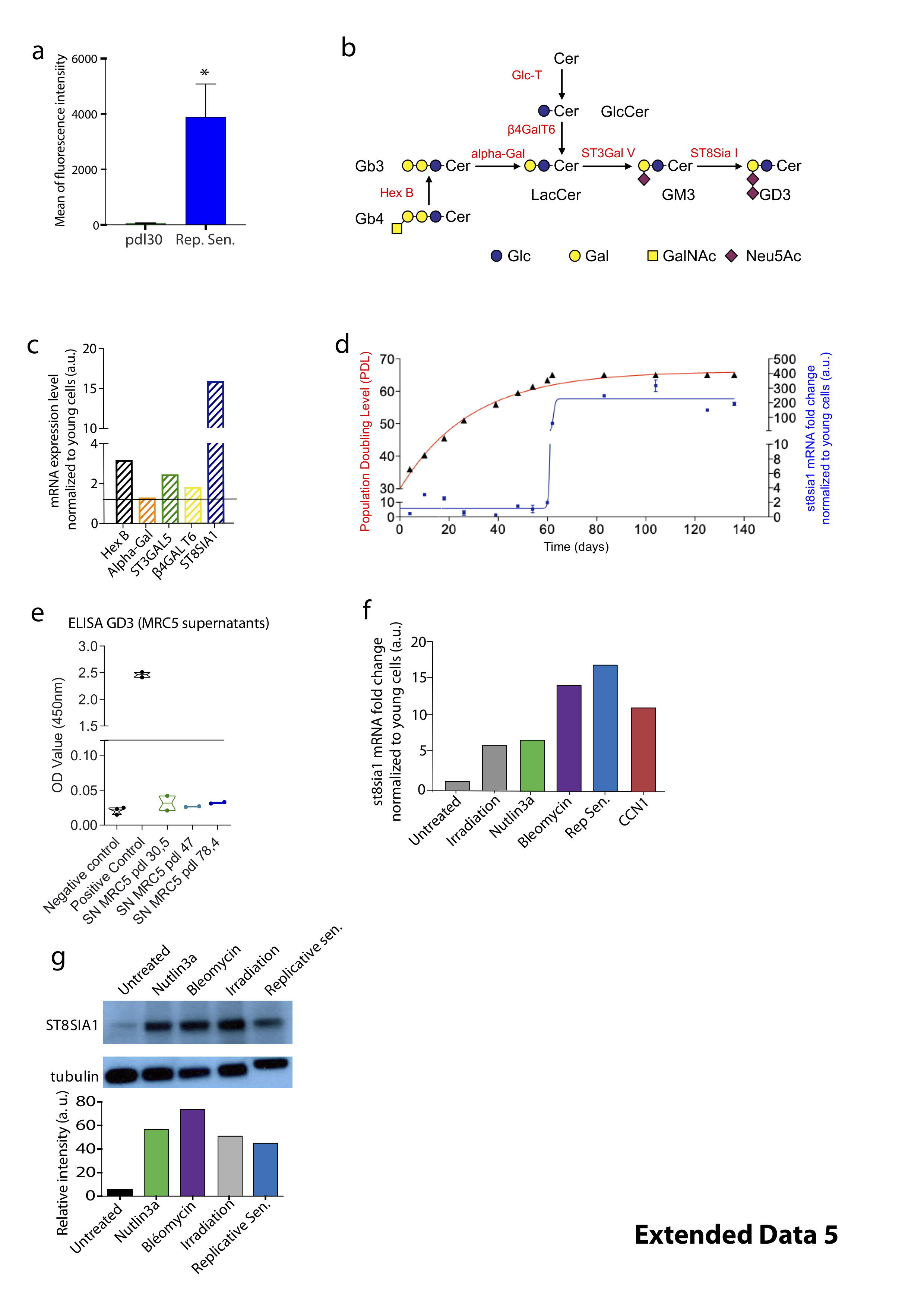

### Extended Data 6

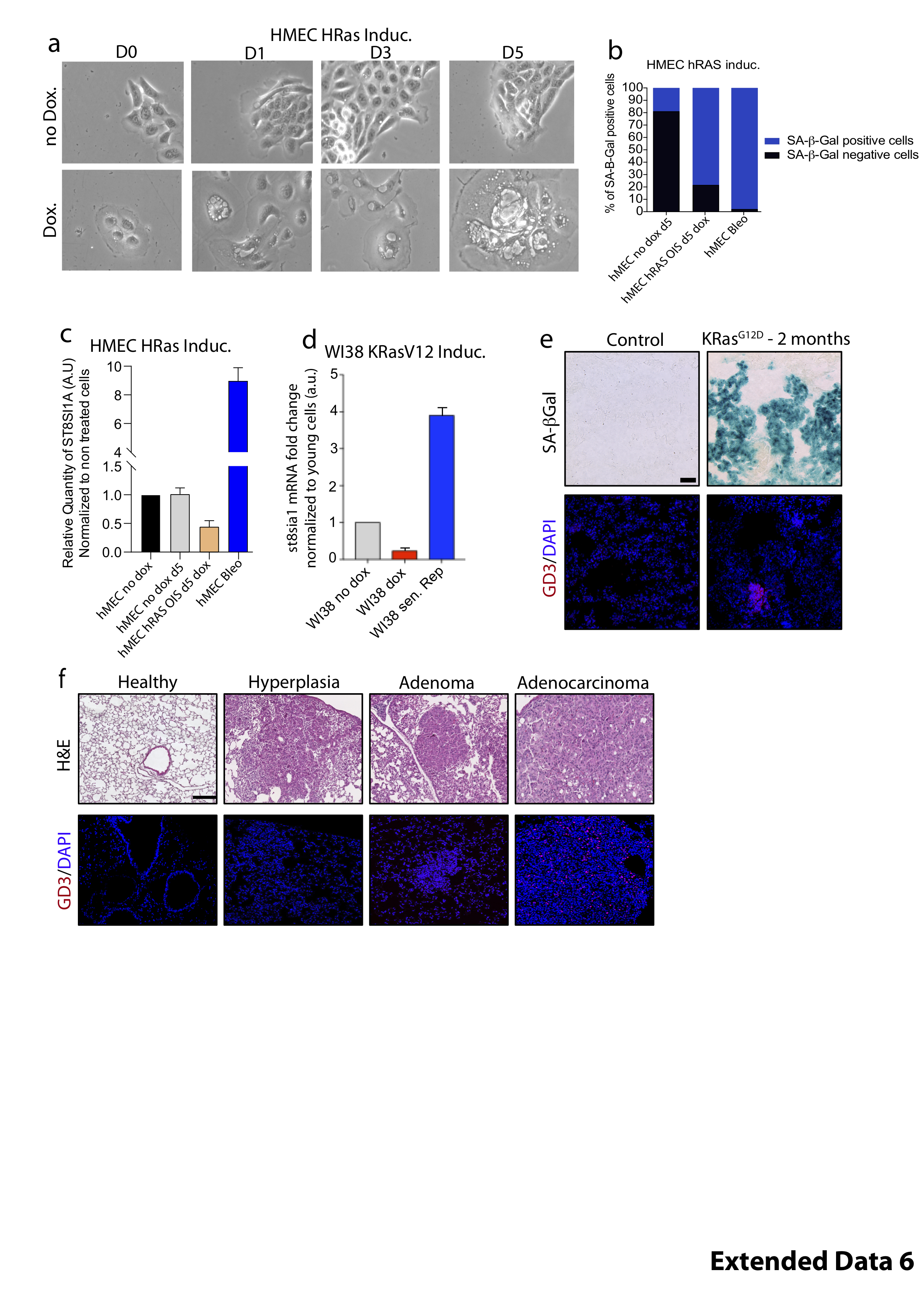

### Extended Data 7

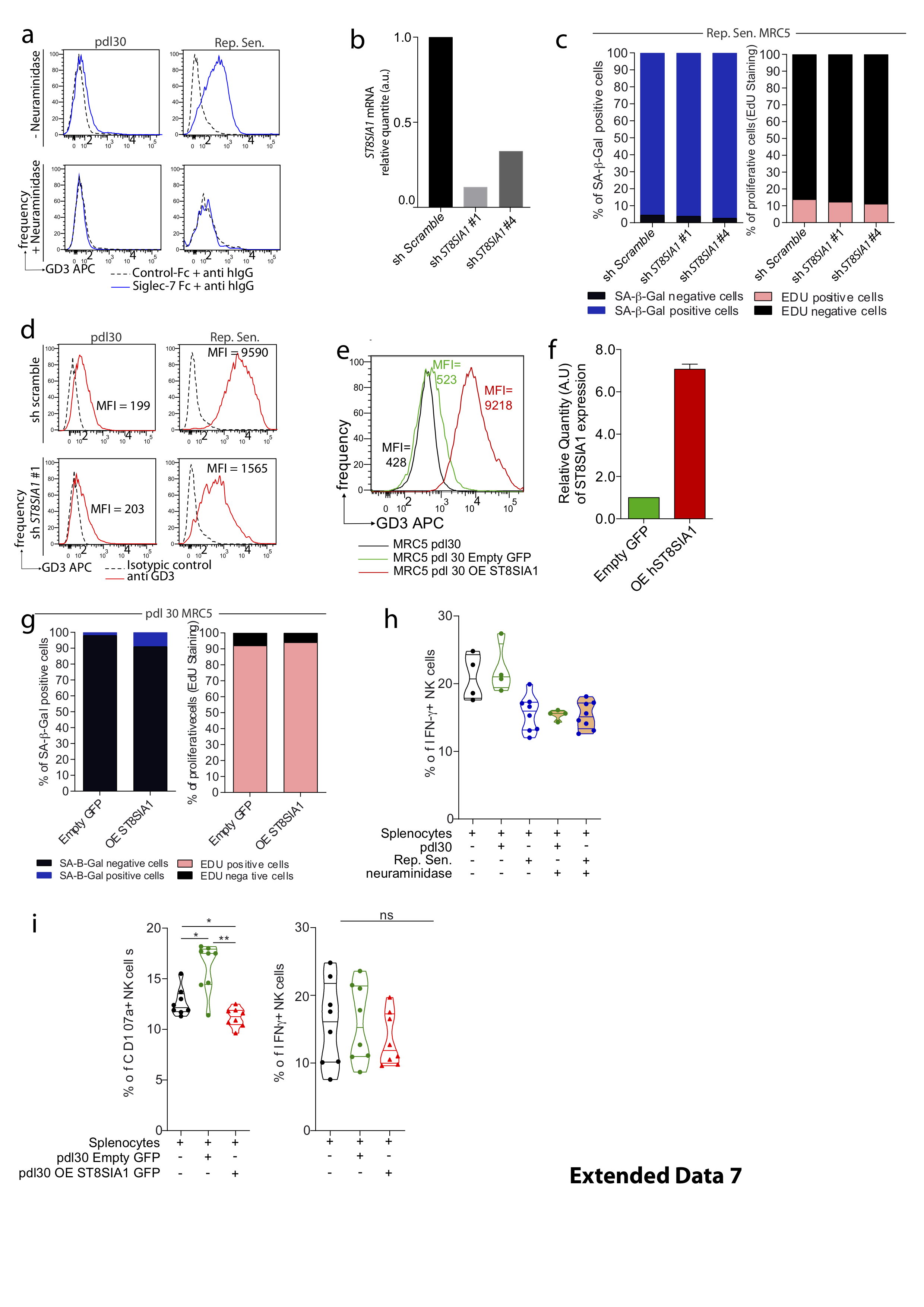

### Extended Data 8

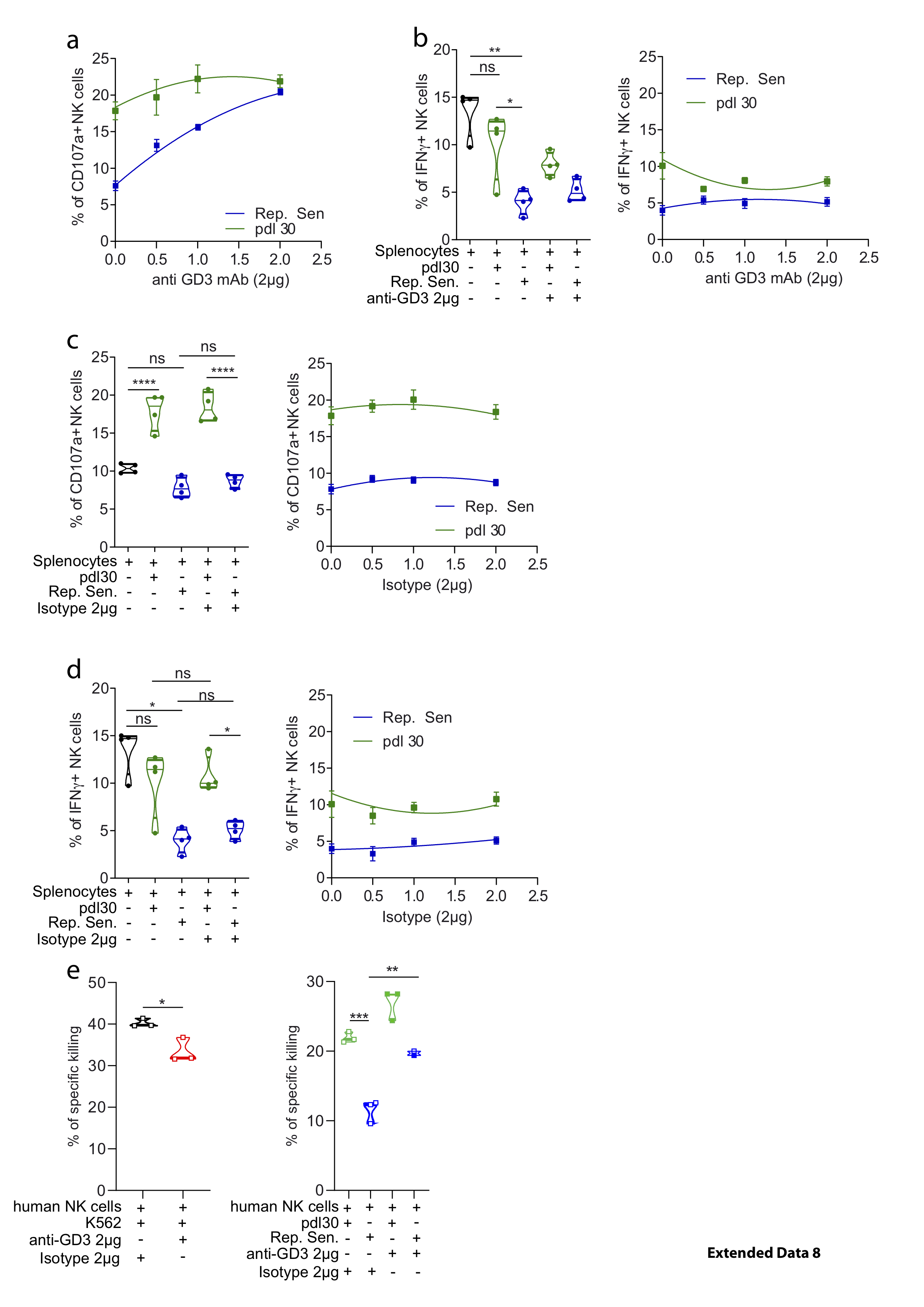

### Extended Data 9

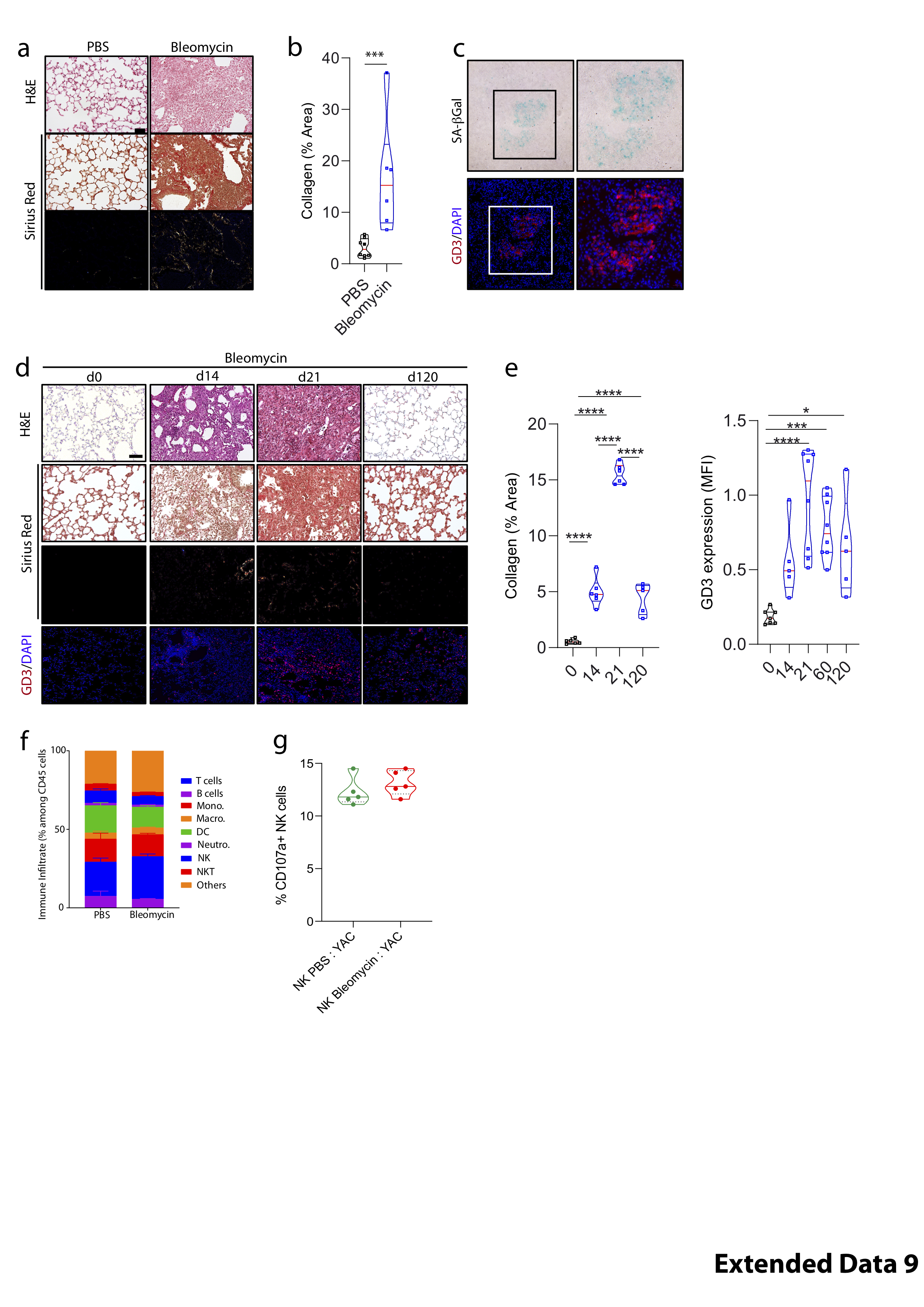

### Extended Data 10

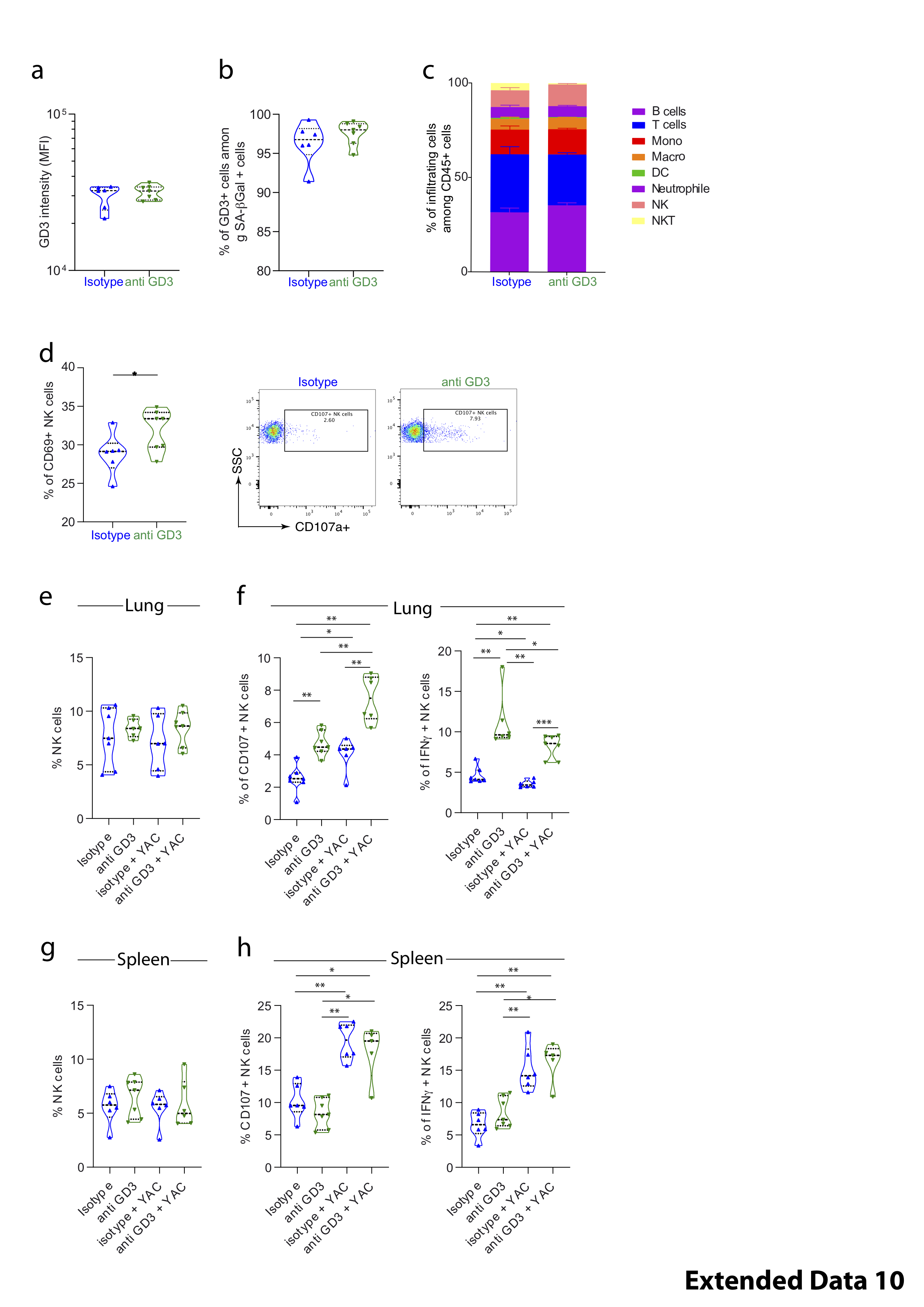

### Extended Data 11

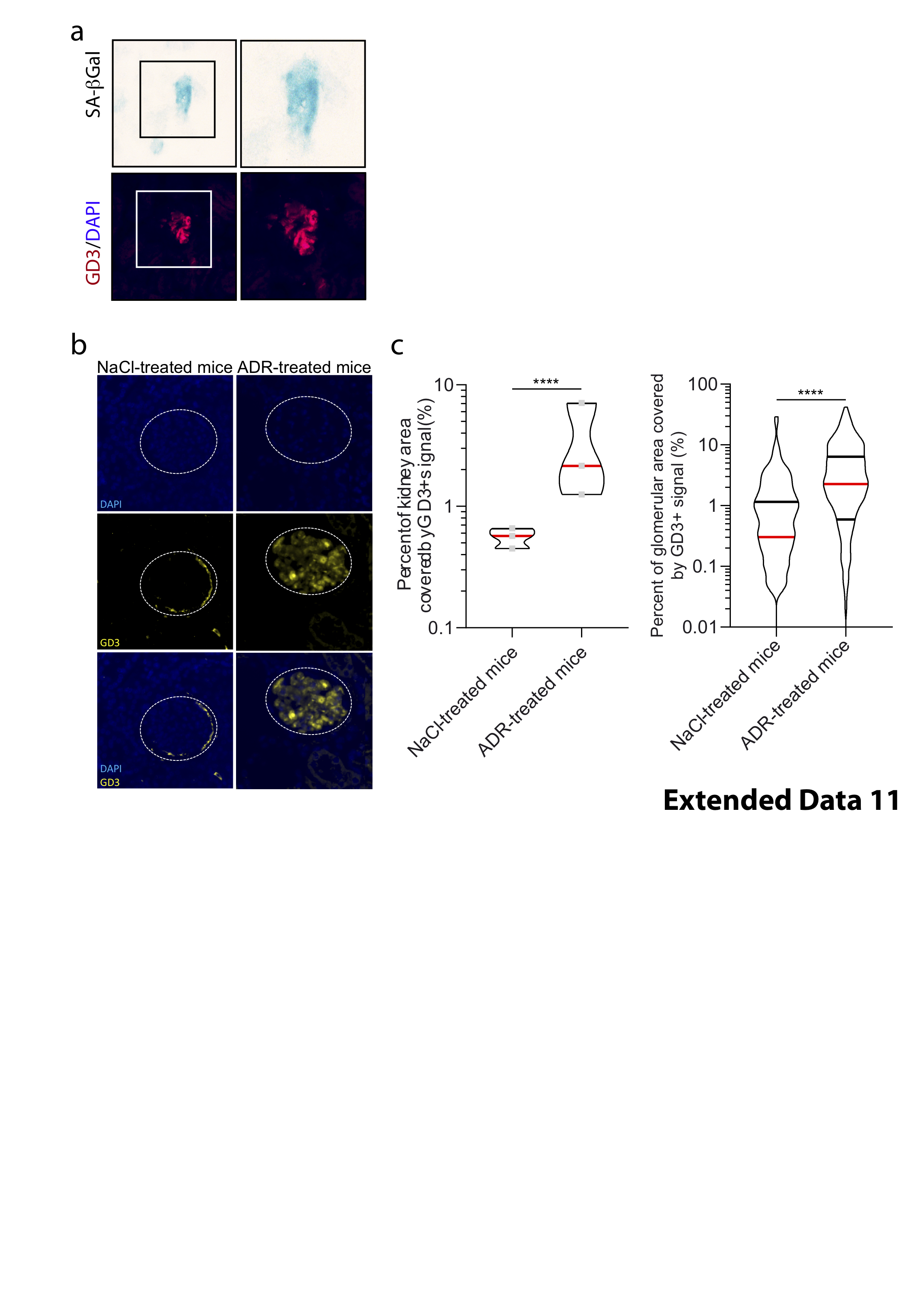

### Extended Data 12

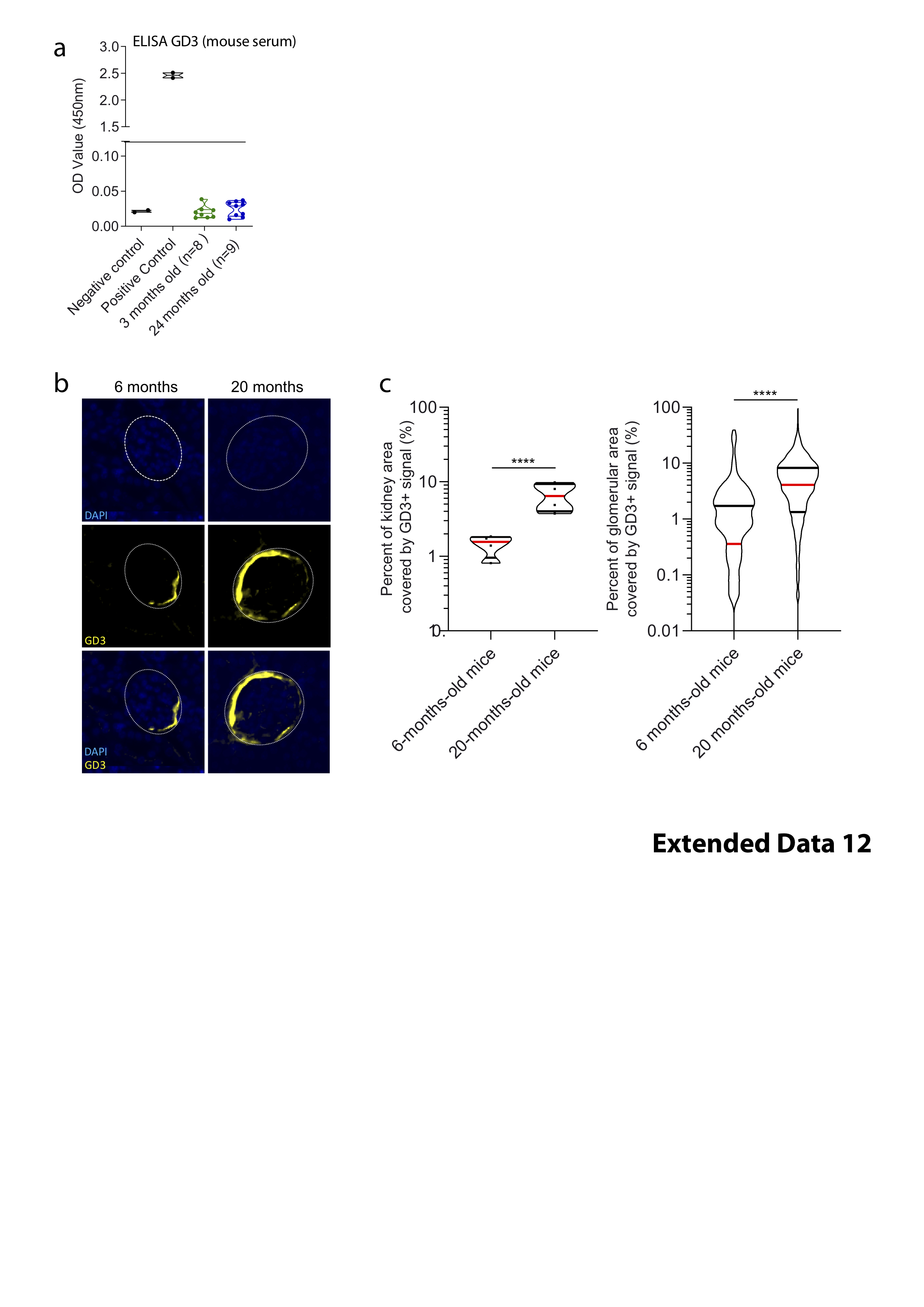

### Extended Data 13

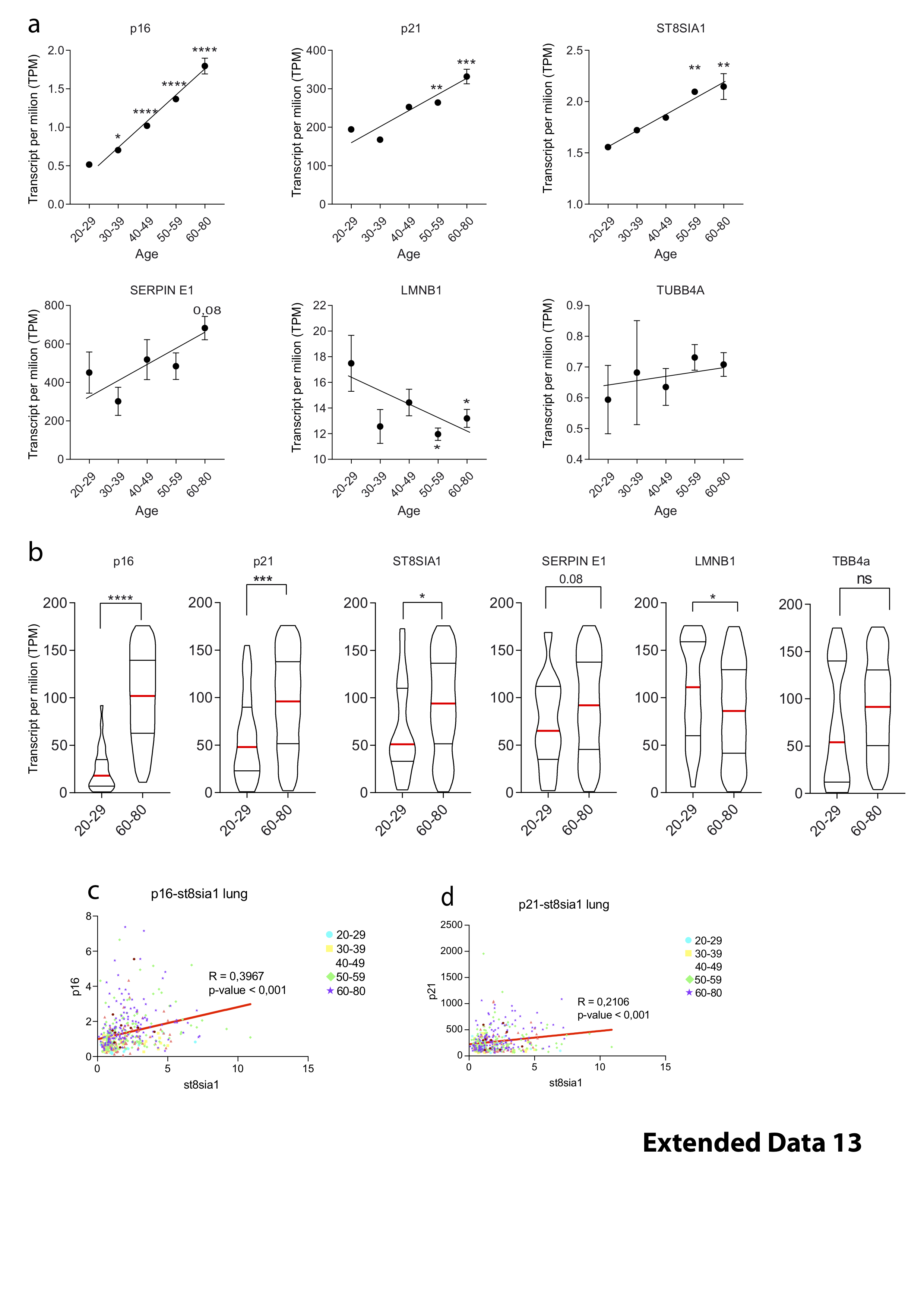
